# Low forces push the maturation of neural precursors into neurons

**DOI:** 10.1101/2022.09.07.507054

**Authors:** Sara De Vincentiis, Matteo Baggiani, Francesca Merighi, Valentina Cappello, Jakub Lopane, Mariachiara Di Caprio, Mario Costa, Marco Mainardi, Marco Onorati, Vittoria Raffa

**Author notes:** Authors equally contributing (Matteo Baggiani, Francesca Merighi).

## Abstract

Mechanical stimulation modulates neural development and neuronal activity. In a previous study, we proposed magnetic “nano-pulling” as a tool to generate active forces. By loading neural cells with magnetic nanoparticles (MNPs), a precise force vector is remotely generated through static magnetic fields. In the present study, human neural stem cells (NSCs) were subjected to a standard differentiation protocol, in the presence or absence of nano-pulling. Under mechanical stimulation, we found an increase in the length of the neural processes which showed an enrichment in microtubules, endoplasmic reticulum, and mitochondria. A stimulation lasting up to 52 days induced a strong remodelling at the level of synapse density and a re-organization of the neuronal network, halving the time required for the maturation of neural precursors into neurons. We then injected the MNP-loaded NSCs into mouse spinal cord slices, demonstrating that nano-pulling stimulates the elongation of the NPC processes and modulates their orientation even in an *ex vivo* model system. To the best of our knowledge, this is the first evidence showing that active mechanical stimuli can guide the outgrowth of NSCs transplanted into the spinal cord tissue. Our findings suggest that MNPs play an important role in neuronal maturation which could be applied in regenerative medicine.

## INTRODUCTION

Central nervous system (CNS) development is regulated by a complex interplay of chemical and mechanical cues, which direct neural progenitor cell (NPC) proliferation, migration, and differentiation through sequentially and spatially coordinated events. The first neural structure that arises in the embryo is the neural tube, which consists of a pseudostratified layer of neuroepithelial cells (Stiles & Jernigan, 2010). These early neural stem cells (NSCs) represent the founders of neurons, astrocytes, and oligodendrocytes (Breunig et al., 2011; Silbereis et al., 2016). Significant advances have been made in clarifying the chemical cues involved in this process (Barkho et al., 2006; Shen et al., 2004; Temple, 2001). However, starting with the formation of the neural tube, many mechanical events mark the major steps of CNS morphogenesis, such as the tube expansion, which then form the three primary encephalic vesicles, neural migration, axon outgrowth, pathfinding, and gyrification of the cerebral cortex (Essen, 1997; Stiles & Jernigan, 2010). The role of mechanical cues is however still poorly understood (Kumar et al., 2017) and is attracting attention (Abuwarda & Pathak, 2020; Luo & O’Leary, 2005; Vittoria Raffa, 2022; Vining & Mooney, 2017). Mechanical stimuli can be classified as passive and active which interplay during neurodevelopment (Discher et al., 2005). Passive mechanical stimuli influence cells through matrix stiffness, porosity, and topography. The mechanical properties of the extracellular matrix change over time and space in the developing neural tissue (Chatelin et al., 2010; Franze, 2013), and coordinate with chemical signaling cascades in building the CNS. Conversely, active stimuli are generated by tissue re-shaping due to cell proliferation, migration, cell-cell, and cell-matrix adhesion. For example in the neural tube or the cortex gyri, they generate and transmit over long distances, pulling and pushing forces that stretch, compress, bend or twist the surrounding cells (Abuwarda & Pathak, 2020; Nishimura et al., 2012; Suzuki et al., 2012).

The effects of passive stimuli on NPC differentiation have been investigated above all using 2D and 3D scaffolds which can be precisely designed with tailored features (Abdel Fattah & Ranga, 2020; Rammensee et al., 2017). Unfortunately, active stimuli are challenging to investigate because of the limited availability of biophysical tools to generate them *in vitro* and *in vivo*. Fire-polished glass pipettes have been used to influence neural stem cell differentiation along a neuronal or astrocytic lineage by modulating the activity of stretch-activated ion channels (Pathak et al., 2014). Stretchable membranes actuated by a linear translation stage have been applied to generate high mechanical tension *in vitro* for inducing NSC differentiation towards mature neuronal cells (Arulmoli et al., 2015; Chang et al., 2013).

Remotely controlled magnetic manipulation is now being used to generate active mechanical stimuli in neural cells (Dai et al., 2019; de Vincentiis et al., 2020; Kunze et al., 2015; Pita-Thomas et al., 2015; Vittoria Raffa et al., 2018; Riggio et al., 2014). The “nano-pulling” paradigm is based on cell loading with magnetic nanoparticles (MNPs) combined with a magnetic field gradient to generate a magnetic force. Since cells behave like a viscoelastic fluid, the magnetic force generated is likely to be dissipated through deformation, and the rigid elements of the cell cytoskeleton thus undergo stretching.

Magnetic nano-pulling allows the *in vitro* chronic stimulation of cells with extremely low forces (<1 nN) and continuous load application for days or weeks, mimicking *in vivo* developmental conditions. This process is named stretch growth (SG) (Pfister et al., 2004) and is accompanied by the addition of new mass and axonal cytoskeleton remodeling (de Vincentiis et al., 2020; Falconieri Alessandro et al., 2022). In developing neurons, nano-pulling is associated with axonal elongation, sprouting, and neuron maturation (de Vincentiis et al., 2020; Falconieri et al., 2021; Falconieri Alessandro et al., 2022; Pita-Thomas et al., 2015; Steketee et al., 2011; Wang et al., 2020). This suggests that SG may also influence neurogenesis. Recently, magnetic nano-pulling was used to promote the neural differentiation of stem cells (Dai et al., 2019). However, many questions remain: *(i)* does the *in vitro* chronic application of extremely low active forces mimicking endogenous *in vivo* conditions influence NSC differentiation? In line with the effects observed in developing neurons, *(ii)* is maturation affected? *(iii)* Are the mechanisms behind SG conserved between different development stages, with similarities in the responses of immature and mature neurons? Finally, *(iv)* can mechanical stimuli induce SG of neurons grafted into a nervous tissue?

To address these points, in this work, spinal cord (SC)-derived human neuroepithelial stem (NES) cells were used (Dell’Anno et al., 2018; Onorati et al., 2016). SC-NES cells show great neurogenic potential, leading to mature neurons with extended complex neurites, as well as astrocytes and oligodendrocytes, thus demonstrating their multipotentiality. In addition, SC-NES cells injected in rodent models of spinal cord lesions can establish functional circuits leading to an amelioration of motor deficits (Dell’Anno et al., 2018), thus showing great potential for the treatment of spinal cord injuries (SCIs).

In this work, SC-NES cells were subjected to a standard differentiation protocol, in the presence or absence of nano-pulling. Nano-pulling was found to induce a strong remodeling at the level of the cytoskeleton, synapses, and neural network, thus reducing the time required for the maturation of neural precursors into mature neurons. The morphological and functional remodeling induced by nano-pulling matches our previous observations in hippocampal neurons (de Vincentiis et al., 2020). This thus suggests that mechanosensitivity is a unified and well-conserved mechanism in development aimed at regulating neurogenesis and neuronal terminal differentiation. Lastly, our results support the hypothesis that nano-pulling can induce SG in the spinal cord tissue, thus opening up fascinating opportunities regarding the use of mechanical stimuli for the treatment of spinal cord injury (SCI).

## RESULTS

### Nano-pulling of SC-NES cells induces stretch-growth

SC-NES cells were differentiated following a previously described protocol (Dell’Anno et al., 2018). Briefly, cells were incubated in the pre-differentiation medium for seven days *in vitro* (DIV) and in the differentiation medium from DIV7 up to DIV90, when they differentiate into mature neurons (Fig. 1A). In order to test the best paradigm for MNP administration, SC-NES cells were loaded with MNPs four hours after the beginning of the pre-differentiation phase (DIV0) or four hours after the beginning of the differentiation phase (DIV7). At DIV8, both groups were exposed to the external magnetic field (“stretch” group) or placed in a mock apparatus producing a null magnetic field (“ctrl” group) for 48 hours. A single-neural process tracing analysis was then performed on TUBB3-positive cells (see supplementary Fig. S1A).

**Figure 1.**
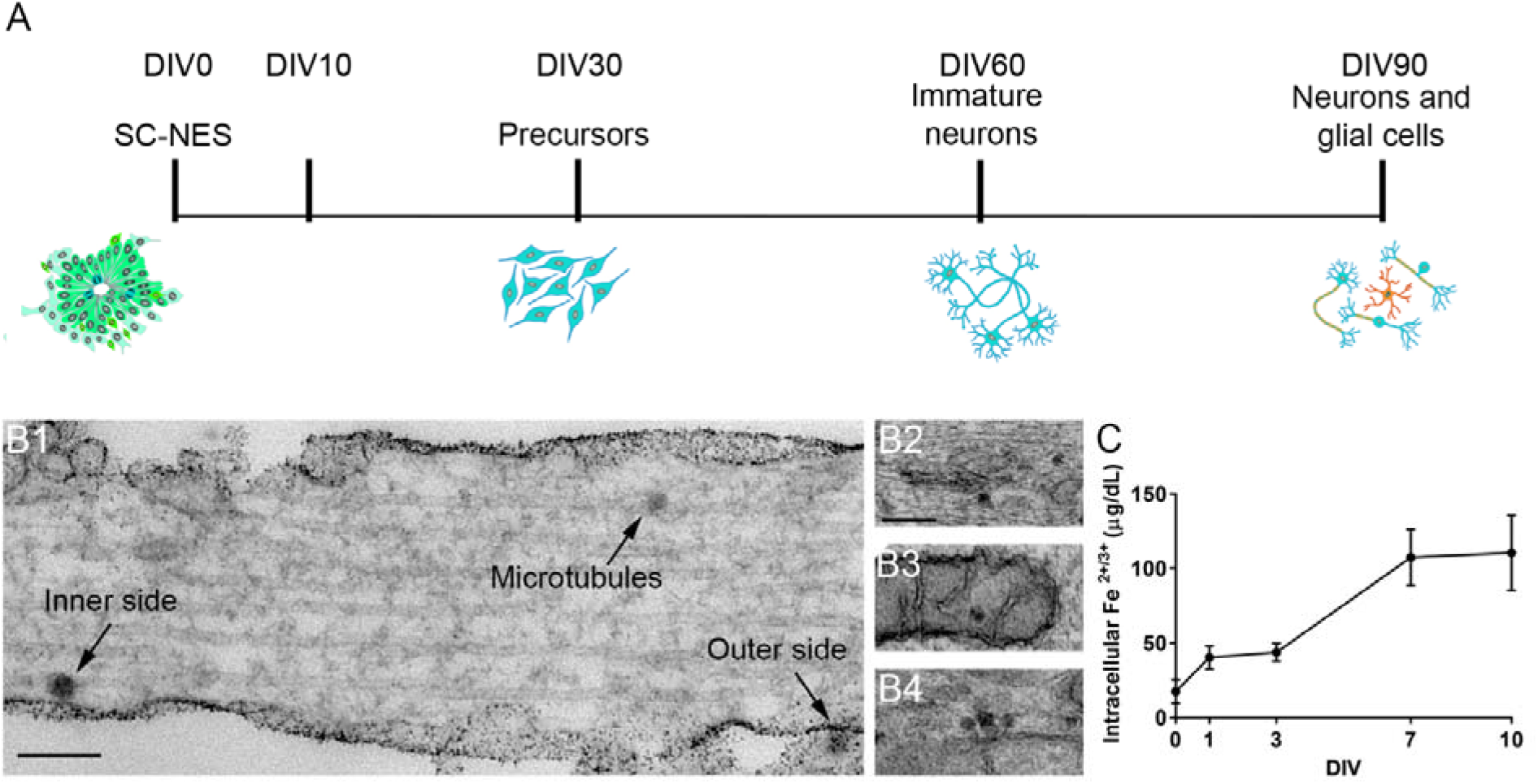
Loading of SC-NES cells with MNPs. A) Schematic representation of the differentiation protocol from undifferentiated SC-NES cells (DIV0) to neurons and glial cells (DIV90). B) Representative TEM micrographs of the localization of MNPs inside a neural process of a NES cell at DIV10. Scale bar 200 nm. B1) MNPs localize both in the outer and inner sides of the cellular membrane, as well as inside the cytoplasm, where, eventually, they are associated with microtubules and endoplasmic reticulum (B2), or inside mitochondria (B3) and a few clusters of 2-3 particles were observed (B4). C) Intracellular level of Fe^2+/3+^ at different time points after MNP loading (0 time point); n=4.

For both conditions, an increase in neural process length in the stretched group was observed but not in the control group (p<0.0001, Fig. S1B). Specifically, the two loading protocols resulted in a length increase of 65.8 ± 3.3% and 75.7 ± 3.8% for DIV0 and DIV7, respectively, which was not statistically different between the two stretched groups (p=0.29, Fig. S1B). This suggests that the time of MNP administration may not be crucial for nano-pulling to exert its effect on neural process outgrowth. For the subsequent experiments, MNPs were added four hours after the beginning of the differentiation phase (DIV7). MNPs alone or the magnetic field alone had no effects on the length of the neural processes (Fig. S1C).

### Nano-pulling generates a pico-Newton force on neural processes

MNP degradation dynamics were characterized at different time points (DIV1, 3, 7, and 10) by measuring the intracellular levels of Fe^2+^ and Fe^3+^. In line with our previous data from mouse hippocampal neurons (de Vincentiis et al., 2020; De Vincentiis et al., 2021), the intracellular iron level dramatically increased after DIV3, marking the beginning of particle degradation. The degradation level then reached a plateau at DIV7, corresponding to a steady state in the dynamics of iron accumulation and elimination (Fig. 1C).

MNP intracellular localization was studied by transmission electron microscopy (TEM) imaging at DIV10. Again in line with our previous results (de Vincentiis et al., 2020; V. Raffa et al., 2018), MNPs were found to be located in the neural processes at both the inner and outer sides of the cell membrane (Fig. 1B1) and diffused within the cytoplasm, associated with microtubules (MTs) or endoplasmic reticulum (ER) and inside the mitochondria (Fig. 1B1-B3).

Clusters of 2-3 particles (Fig. 1B4) were rarely observed, indicating that MNPs are internalized as single particles. The number of MNPs was quantified in TEM images and normalized per unit of the neural process volume analyzed (1.72 ± 0.31 MNPs·μm^−3^, n= 30). The neural process was then modeled as a cylinder and the mean volume was calculated based on the mean length and caliber of control neural processes reported in the next section. Specifically, the mean total volume of MNPs in the neural process was found to be 4.24 ± 0.75 μm^3^. Using Eq. 1 (see Materials and Methods), the mean force generated on-axis in the neural process was estimated to be 10.55 ± 1.85 pN.

### Responsiveness of SC-NES cells to SG is cell-stage independent

To test the responsiveness of SC-NES cells to SG during the differentiation process and to collect quantitative data, MNPs were added at different time points during terminal differentiation (DIV7, DIV27, DIV57, and DIV87) and cells were stimulated for the subsequent 48 hours (Fig. 2A). The SG of stretched samples was observed at all the time points tested (p<0.0001, Fig. 2A1-A4), demonstrating that cells respond to stretching throughout the process of cell differentiation. However, the highest increase in neural process length was observed at DIV10 (65.8 ± 5.9% at DIV10, 33.2 ± 4.3% at DIV30, 41.3 ± 3.4% at DIV60, and 53.5 ± 4.2% at DIV90; Fig. 2C). The increase in neural process length between the control and stretched conditions at DIV10 is easily detectable when cells are cultured in microfluidic devices. Specifically, the microfluidic system is composed of two chambers connected by microfluidic channels: cells are seeded in one compartment and neural processes can easily cross the channels and invade the opposite chamber. Under the effect of directional nano-pulling, stretched neural processes show a strong tendency to reach longer distances than the unstretched group (Fig. 2B).

**Figure 2.**
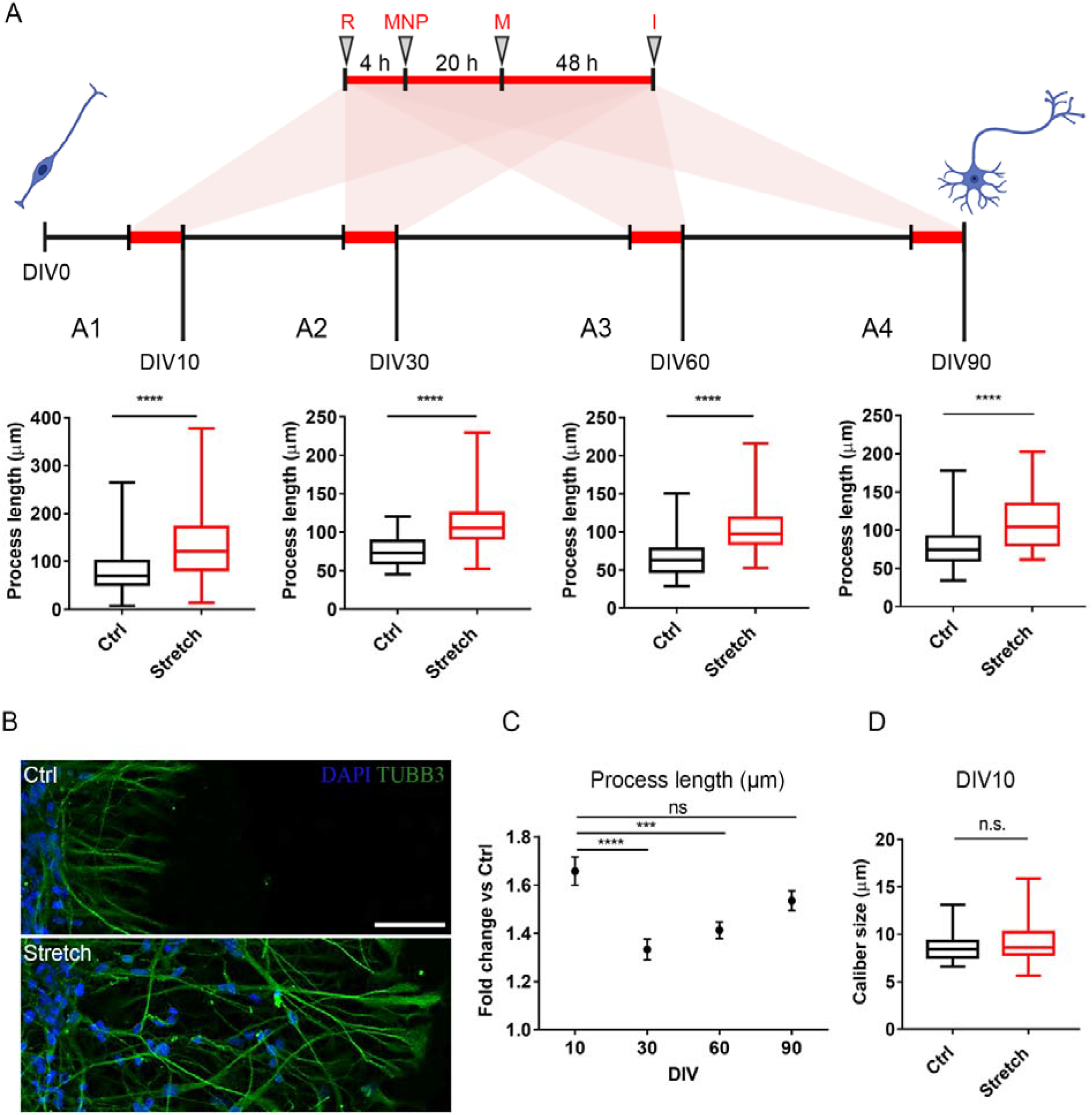
Short-term stretching assay. (A) Schematic representation of the short-term experimental design during the differentiation of SC-NES cells into neurons. At DIV7, 27, 57, and 87 cells are replated (R), 4 h later MNPs are added, and 20 h later the “stretch” groups are exposed to the magnetic field (M). After 48 h of continuous stretching, samples are fixed and subjected to imaging (I). (A1) Neural process length at DIV10, p<0.0001. (A2) Neural process length at DIV30, p<0.0001. (A3) Neural process length at DIV60, p<0.0001. (A4) Neural process length at DIV90, p<0.0001. (A1-A4) Box plot, min-to-max, n=400 processes. Mann-Whitney test. (B) Representative images of TUBB3 (green) and DAPI (blue) in control (upper part) and stretched (bottom part) conditions of NES cells cultured in microfluidic devices at DIV10. Scale bar 100 μm. (C) Fold change comparison of stretched versus control conditions at different DIV. Mean ± SEM, n=400 processes. (D) Neural process caliber at DIV10. Box plot, min-to-max, n=40 processes. Mann-Whitney test, p=0.15.

These results were not related to the specific NES cell line, since a similar outcome was observed in a different NES cell line derived from human-induced pluripotent stem (NES-iPS) cells (Fig. S2).

### Nano-pulling induces mass addition and accumulation of microtubules, mitochondria, and endoplasmic reticulum in neural processes

To demonstrate that the lengthening of the neural processes is an example of real growth rather than merely viscoelastic deformation, their caliber was estimated under the maximum elongation rate *(i.e*., DIV 10, Fig. 2D). No statistically significant difference was detectable between the control and stretched groups (p = 0.15), suggesting that mass addition may occur.

This prompted us to investigate the cellular mechanisms underlying this phenomenon by comparing the ultrastructure of control and stretched neural processes. TEM imaging was used to trace MTs and to estimate the linear density as the ratio of their number in a longitudinal cross-section to the neural process caliber (Fig. 3A1). A strong increase in MT linear density was found (p<0.0001, Fig. 3A2). Specifically, 2.8 ± 0.1 and 4.4 ± 0.1 MTs per μm were measured in control and stretched conditions respectively, with a net increase of approximately 60%. Similarly, enrichment of ER cisternae was revealed in stretched samples by analyzing TEM images (ER, Fig. 3A1), which showed an overproduction of ER cisternae in stretched neural processes (80.6 ± 14.7%, p<0.0001, Fig. 3A3). The accumulation of ER was confirmed by immunostaining for KDEL (an increase of 71.8 ± 6.6 % in the stretch group versus the control group, p<0.0001, Fig. 3B2). Considering the role played by the ER in local lipid and protein production (Valenzuela & Perez, 2015), these results seem to support the hypothesis that a new mass could be added locally in response to SG.

**Figure 3.**
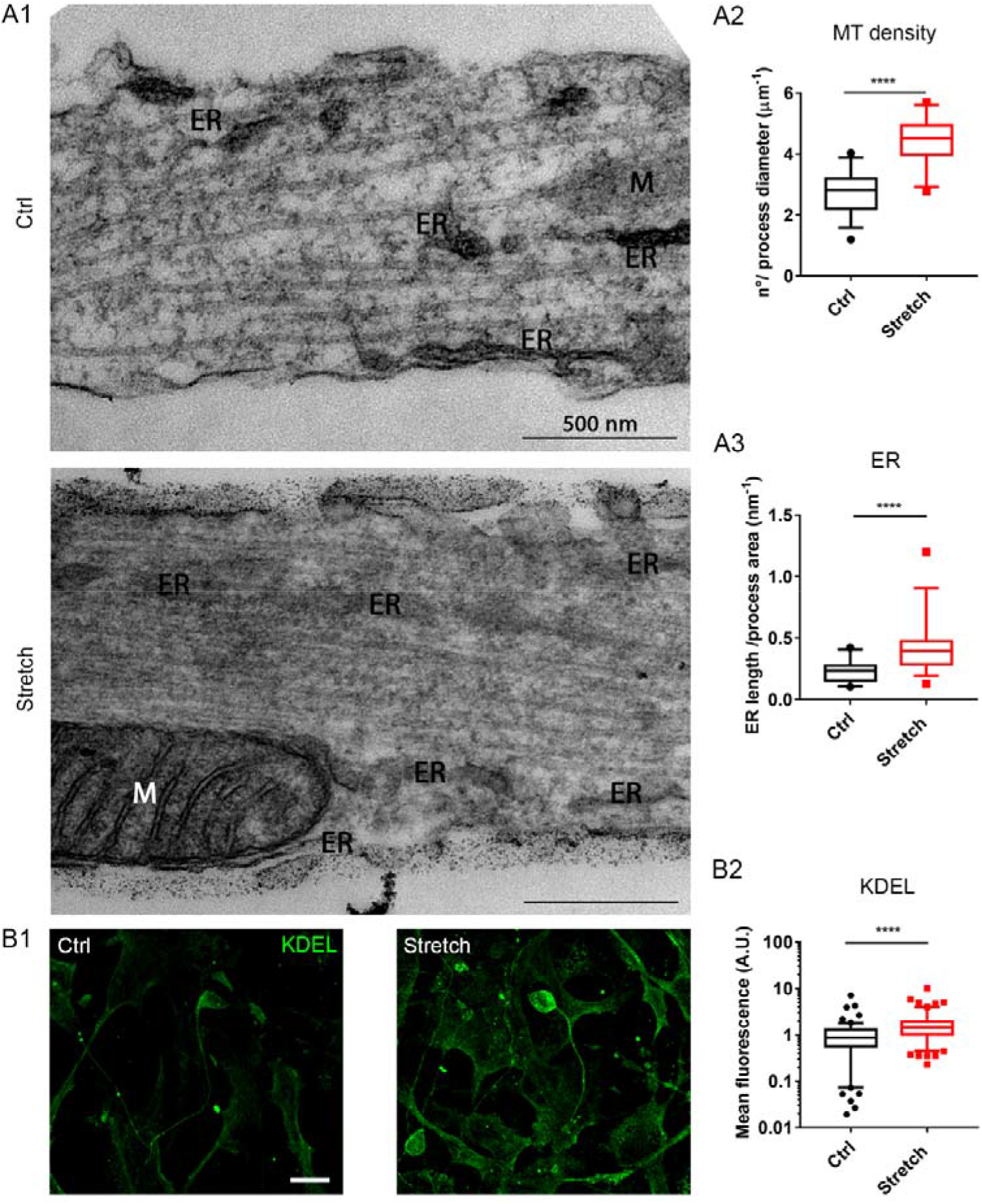
Accumulation of mitochondria, microtubules and endoplasmic reticulum in stretched SC-NES cells. A1) Representative micrographs of SC-NES cells at DIV10 in control and stretched conditions. Mitochondria (M) and endoplasmic reticulum (ER) highlighted. Scale bar 500 nm. A2) Quantification of microtubule linear density. Box plot, 5-95 percentile, n=35 processes. T-test for unpaired data, p<0.0001, t=9.496, df=68. A3) Quantification of ER cisternae. Box plot, 5-96 percentile, n=30 processes. Mann- Whitney test, p<0.0001. B1) Representative pictures of SC-NES cells at DIV10 in control and stretched conditions stained with KDEL (green). Scale bar 10 μm. B2) Quantification of KDEL fluorescence. Box plot, 5-95 percentile, n=120 processes. Mann- Whitney test, p<0.0001.

### Nano-pulling shapes cell networking

Previous studies analyzing the effect of active force application have suggested that mechanical exogenous forces may act on NSC commitment, by enhancing the elongation and maturation of NSC-derived neurons (Arulmoli et al., 2015; Chang et al., 2013; Dai et al., 2019). Unlike previous reports, here, SC-NES cells were stimulated from the beginning to the end of the differentiation process, in order to fully describe the effects of SG at different time-points. Specifically, cells were exposed to the stimulus for 8, 22, or 52 days after the beginning of the differentiation protocol (Fig. 4A). Cells formed a complex network with interconnected neural processes (Fig. 4B). The extension of neural processes for each soma was estimated as the ratio of the TUBB3-positive area by the number of somata in the region of interest (ROI).

**Figure 4.**
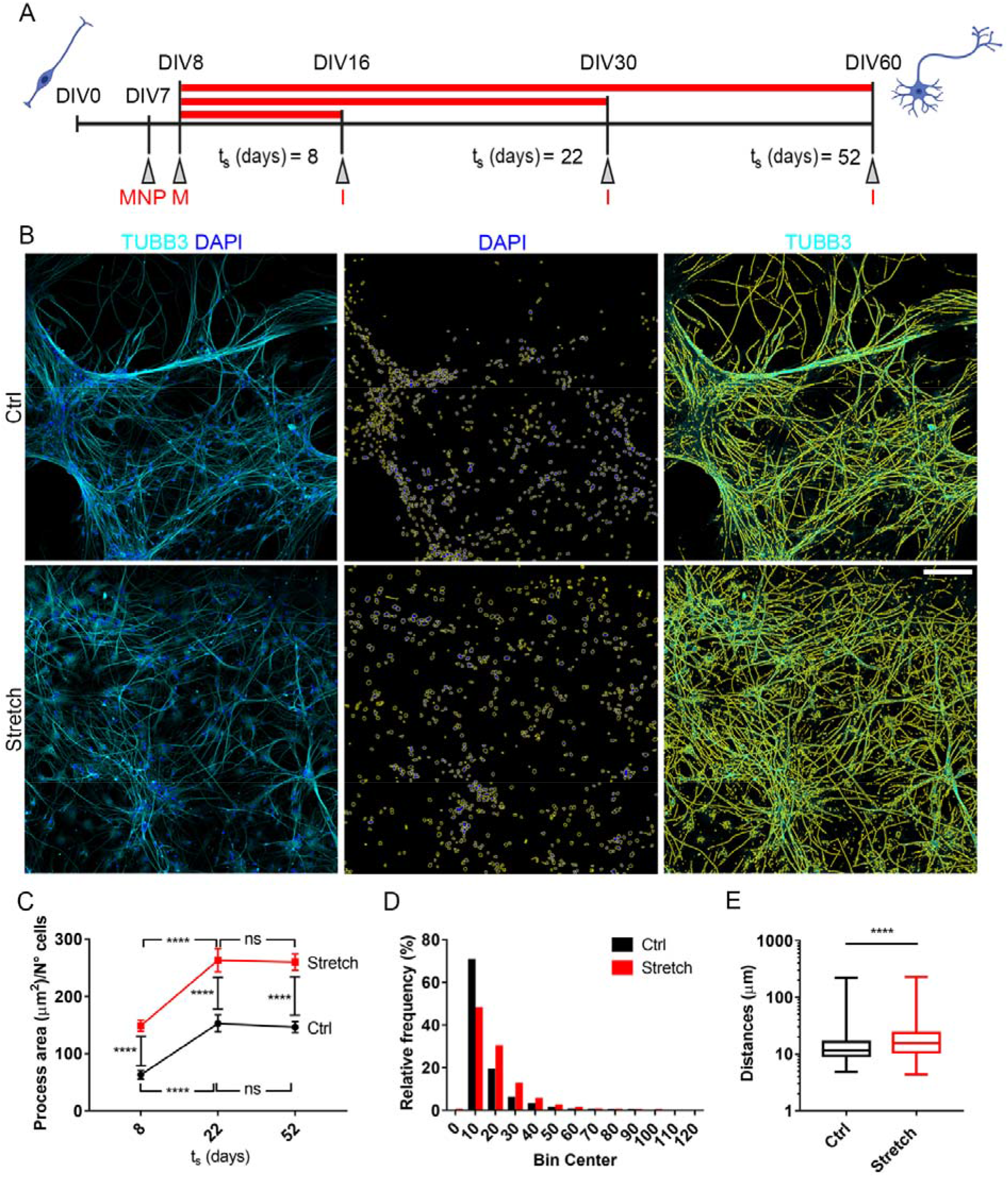
Long-term stretching assay. (A) Schematic representation of the long-term experimental design. At DIV7 MNPs are added and 20 h later the “stretch” groups are placed inside a magnetic applicator (M). After 8, 22, and 52 days of stretching, samples are fixed and subjected to imaging (1). (B) Representative images of TUBB3 (cyan) and DAPI (blue) in NES cells at DIV60 in the control and after 52 days of stretching. In yellow are the regions of interest (ROI) of the occupied area. Scale bar 100 μm. (C) Comparison between stretched and control conditions at different stretching times. Mean ± SEM, n=20 pictures for each time point. Two-way ANOVA test with Holm-Sidak *post hoc’s* test. Row factor (Ctrl vs Stretch): p<0.0001, F=37.17. Columns factor (stretching time): p<0.0001, F=89.53. (Dl Relative frequency distribution of the distances between nuclei at DIV60 (t_s_=52 days). n>13474 nuclei. Kolmogorov-Smirnov test, p<0.0001. (E) Analysis of nuclei dispersion. Mean± SEM, n>13474 nuclei. Mann-Whitney test, p<0.0001.

Our data showed a statistically significant increase in the area occupied by neural processes per number of somata in the stretched samples in comparison with the controls (p<0.0001, with an increase of 135.8 ± 6.8%, 72.4 ± 3.6% and 77.5 ± 3.9% after 8, 22, and 52 days of stretching, respectively, Fig. 4C). Cells thus responded to continuous stimulation, even after prolonged exposure and irrespectively of their maturation stage. Interestingly, the elongation curves of control and stretched samples followed essentially the same trend (the curve shifts up) (Fig. 4C). At the earliest time-point (DIV10), stretched samples showed a higher outgrowth rate than the controls (18.75 and 6.25 μm·day^−1^, respectively), but both groups reached a plateau at DIV30, with a null rate, likely due to the interconnection of the neural processes in a complex network formation (for both groups, p<0.0001 between DIV10 and DIV30, and p=0.94 and p=0.98 between DIV30 and DIV60, for control and stretched groups respectively, Fig. 4C).

This raises the question as to whether interconnected neural processes can still respond to SG. At the last time-point (DIV60), the network morphology appeared to be considerably different between the two conditions (Fig. 4B). It is well known that neuronal network morphology is strongly coupled with cellular connectivity (Chklovskii, 2004) and that this kind of organization can occur in response to external stimuli (Pascual-Leone et al., 2005). This aspect was thus further investigated. By employing nuclei dispersion index 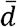 as an estimate of network morphology (Eq. 2, see Material&Methods), a different data distribution was found (p<0.001, Fig. 4D). A statistically significant increase was detected in nuclei dispersion of 28.7 ± 0.2% in stretched samples compared to controls (p<0.0001, Fig. 4E). This finding indicates that nano-pulling can alter the cell network formation.

### Nano-pulling stimulates neuronal differentiation

The data collected raised the question as to whether the effects of the nano-pulling on network re-organization could also impact on neuronal differentiation processes. The cytoskeleton is subjected to changes during neurogenesis in cultured cells (Haendel et al., 1996), with early-stage cells containing only microfilaments, and MTs starting to appear at later stages. The cytoskeleton composition was thus evaluated through TEM analysis after seven days of stretching (DIV10, Fig. 5A). An increase in the number of neural processes containing MTs was observed in stretched samples (ctrl: 46% of neural processes with microfilaments, 54% with microtubules; stretch: 20% with microfilaments, 80% with microtubules; p=0.0001, Fig. 5A5), supporting the idea that nano-pulling induces early remodeling of the cytoskeleton and speeds up the onset of neuronal differentiation.

To further confirm this result, the expression of SOX2 and RBFOX3 was studied at DIV30 and DIV60, *i.e*., 22 and 52 days of stimulation, respectively. SOX2 is a crucial transcription factor for the maintenance of stem cell stemness whose expression wanes and eventually disappears during the completion of differentiation (Zhang, 2014). RBFOX3 (also known as NeuN) is a common marker of mature neurons. After 22 days of stimulation (DIV30), no differences in the percentage of SOX2 positive cells were observed between control and stretch groups (ctrl 81.5 ± 17.4%, stretch 84.4 ± 15.2%; p=0.92, Fig. 5B5). This suggests that cells are still in an undifferentiated stage in both conditions, in keeping with the low levels of RBFOX3-positive cells in both groups (Fig. 5C3). After 52 days of stretching (DIV60), SOX2 staining still failed to show significant differences between the control and stretch groups (p=0.99, Fig. 5B5). Strikingly, at DIV60 the stretch group showed a statistically significant increase in the number of cells positive to RBFOX3, with approximately 31.1 ± 3.9% of positive cells compared to 18.7 ± 1.7% of the control group (p=0.0008, Fig. 5C5).

**Figure 5.**
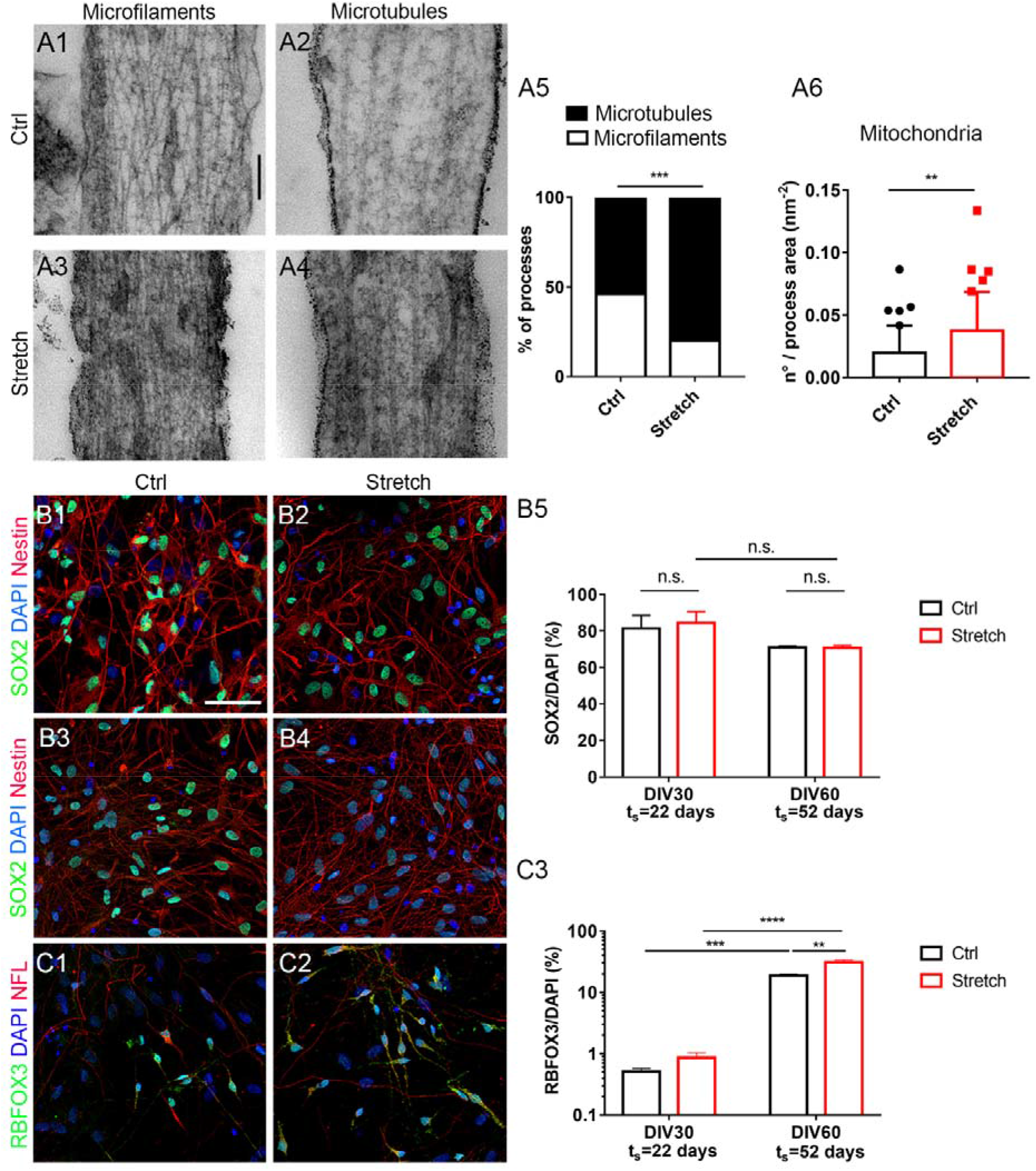
Effect of low forces on the differentiation of SC-NES cells into neurons. (A1-A4) Representative micrographs of microfilaments and microtubules in control and stretched samples at DIV10 (A1-A2 and A3-A4, respectively). A5) Quantification of the cytoskeletal composition. Fisher’s test, n=50 neural processes, p=0.0001. A6) Quantification of mitochondria’s number. Box plot, 5-95 percentile, n=100 processes. Mann-Whitney test, p=0.006. (B1-B4) Representative pictures of SOX2 (green), DAPI (blue), and Nestin (red) immunofluorescence in control (ctrl) and stretched (stretch) cells at DIV30 (B1-B2, respectively), and at DIV60 (B3-B4, respectively). Scale bar: 50 μm. B5) Analysis of SOX2- positive cells at DIV30. Mean ± SEM, n>1000 cells. Two-way ANOVA with Tuckey’s HSD *post hoc* test. Row Main factor 1 (ctrl vs stretch): p=0.10, F=3.07. Main Columns factor 2 (days of stretching): p=0.85, F=0.04. (C1-C2) Representative pictures images of RBFOX3 (green), DAPI (blue), and neurofilament (NFL, red) immunofluorescence at DIV60 in control and stretched conditions. C3) Analysis of RBFOX3-positive cells. Mean ± SEM, n>1000 cells. Two-way ANOVA with Tuckey’s HSD *post hoc* test. Row factor (ctrl vs stretch): p<0.0001, F=453.7. Columns factor (days of stretching): p=0.0014, F=31.19.

In this regard, the mitochondrial pool is known to support neuron differentiation (Iwata & Vanderhaeghen, 2021; Khacho et al., 2019). Specifically, variations in the mitochondrial pool correlate with the change in metabolism (from glycolytic to oxidative) that occurs during neuronal differentiation (Mandal et al., 2011). Here, TEM analysis showed that the number of mitochondria was significantly higher in the stretch group with respect to the control (p=0.006, Fig. 5A6).

Altogether, TEM and confocal microscopy data suggest that SG may push the differentiation of NES cells towards a neuronal phenotype.

### Nano-pulling stimulates functional neuronal maturation

Data shown in Fig. 5 raised the question as to whether long-term stimulation for up to 52 days could enhance the neuronal maturation process.

First, the mean fluorescence intensity of synaptophysin immunoreactivity was measured in the control and stretched neural processes at different time points during the differentiation process (Fig. 6A). The expression of this presynaptic protein was analyzed in order to evaluate the maturation level of the neuronal progeny (Dell’Anno et al., 2018; Kalyani et al., 1998) The temporal window of observation spanned from DIV16, when synapses are still immature, to DIV60, which is characterized by advanced synaptic maturation (Dell’Anno et al., 2018). A statistically significant increase in synaptophysin expression was observed in stretched samples after 8 days (p=0.0003, Fig. 6B) and 22 days (p<0.0001, Fig. 6C) of stimulation compared to the control cultures. On the other hand, at later stages during differentiation, after 52 days of stimulation (DIV60), no significant differences were detected between the stretch and control groups (p=0.91, Fig. 6D). A possible interpretation of these findings is that mechanical nano-pulling may simply speed up the maturation process.

**Figure 6.**
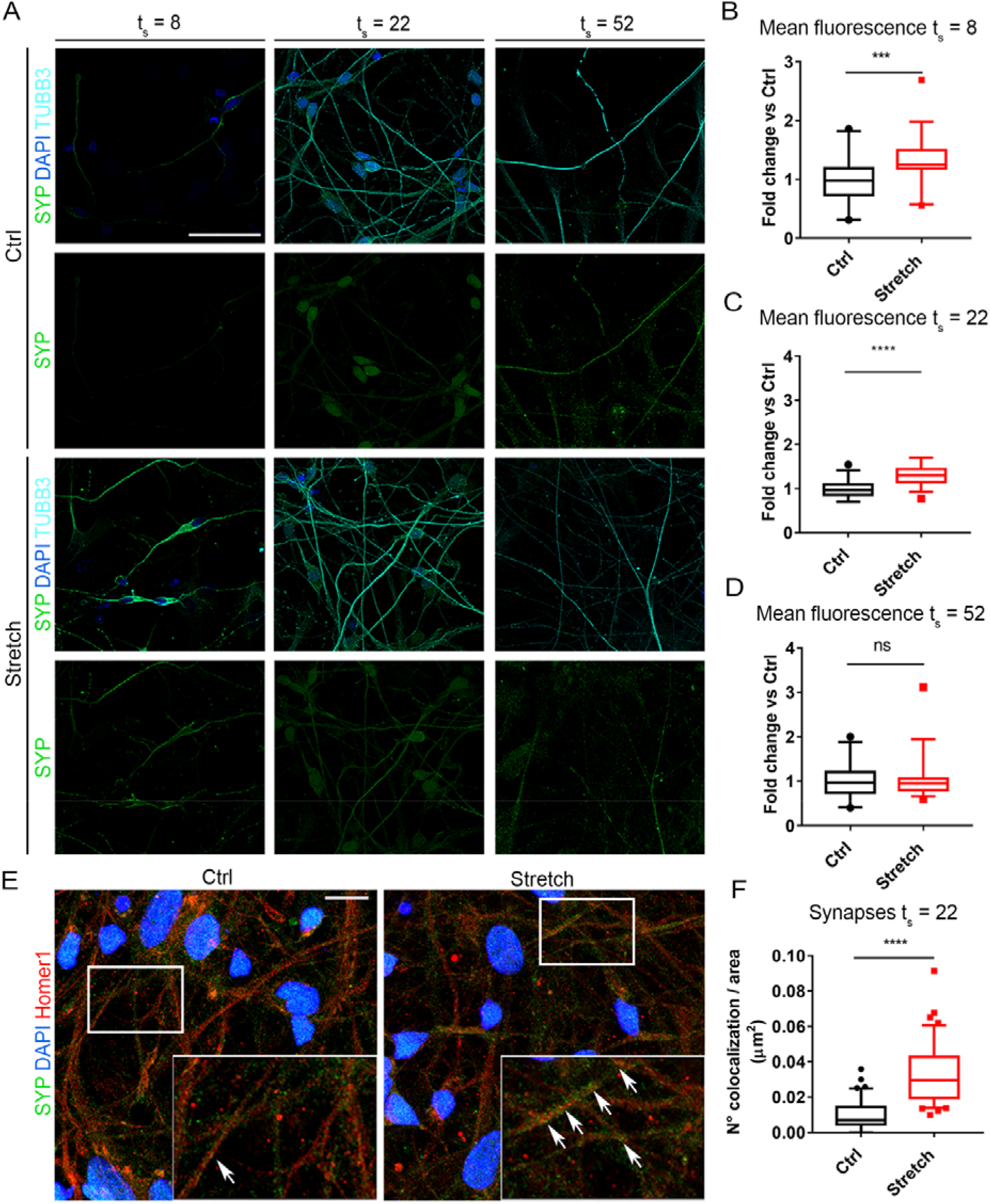
Effects of low forces on neuronal maturation. (A) Representative pictures of Synaptophysin (SVP, green), DAPI (blue), and Tubulin B3 (TUBB3, cyan) in control and stretched conditions at t_s_=8 (DIV16), t_s_=22 (DIV30), and t_s_=52 (DIV60). Scale bar 50 μm. (B) Analysis of SVP mean fluorescence in neural processes at t_s_=8, Mann-Whitney test, p=0.0003. (C) Analysis of SVP mean fluorescence in neural processes at t_s_=22, t-test for unpaired data, p<0.0001, t=6.22, df=70. (D) Analysis of SVP mean fluorescence in neural processes at t_s_=52, Mann-Whitney test, p=0.91. (B-D) Box plot, 5-95 percentile, n=36 pictures. (E) Representative pictures of SVP (green), DAPI (blue), and Homer1 (red) in control and stretched conditions at t_s_=22. White arrows point to the colocalization spots of SVP and Homer1. Scale bar 20 μm. (F) Quantification of the colocalization. Box plot, 10-90 percentile, n=40. Mann-Whitney test, p<0.0001. t_s_: stretching time.

To confirm this hypothesis, colocalization studies between a pre-synaptic and a post-synaptic marker were performed to estimate synapse density following (Verstraelen et al., 2018, 2020). The colocalization between the presynaptic marker synaptophysin and the postsynaptic marker Homer1 was evaluated after 22 days of stretching (DIV30, Fig. 6E). This time point was chosen because *(i)* the difference between stretch and control groups showed the highest significance (p<0.0001, Fig. 6C), and *(ii)* control cultures are expected to be in an immature stage (Fig. 5B3) (Piper et al., 2001). A significant increase was observed in puncta showing apposition between synaptophysin and Homer1 spots, which reflects an increase in fully formed synaptic density, in stretched samples compared to the controls (p<0.0001, Fig. 6F).

To obtain functional evidence of the maturation boost induced by nano-pulling, patch-clamp recordings were employed to assess cellular excitability upon cell depolarization via positive current injection. At DIV30, no action potentials were elicited. In DIV60 cultures, stretched cells fired a higher number of action potentials than the control cells, while at DIV90 no difference was detected between the two groups (Fig. 7A-B). Spontaneous activity in the absence of membrane depolarization was then recorded. At DIV60, 30.0% of stretched cells and 18.2% of control cells showed spontaneous action potentials (Fig. 7C). Interestingly, at DIV90, 59.3% of stretched cells showed spontaneous action potentials, in comparison to 30.8% of control cells (Fig. 7C). On the other hand, no significant differences in resting membrane potential were observed (Fig. 7D).

**Figure 7.**
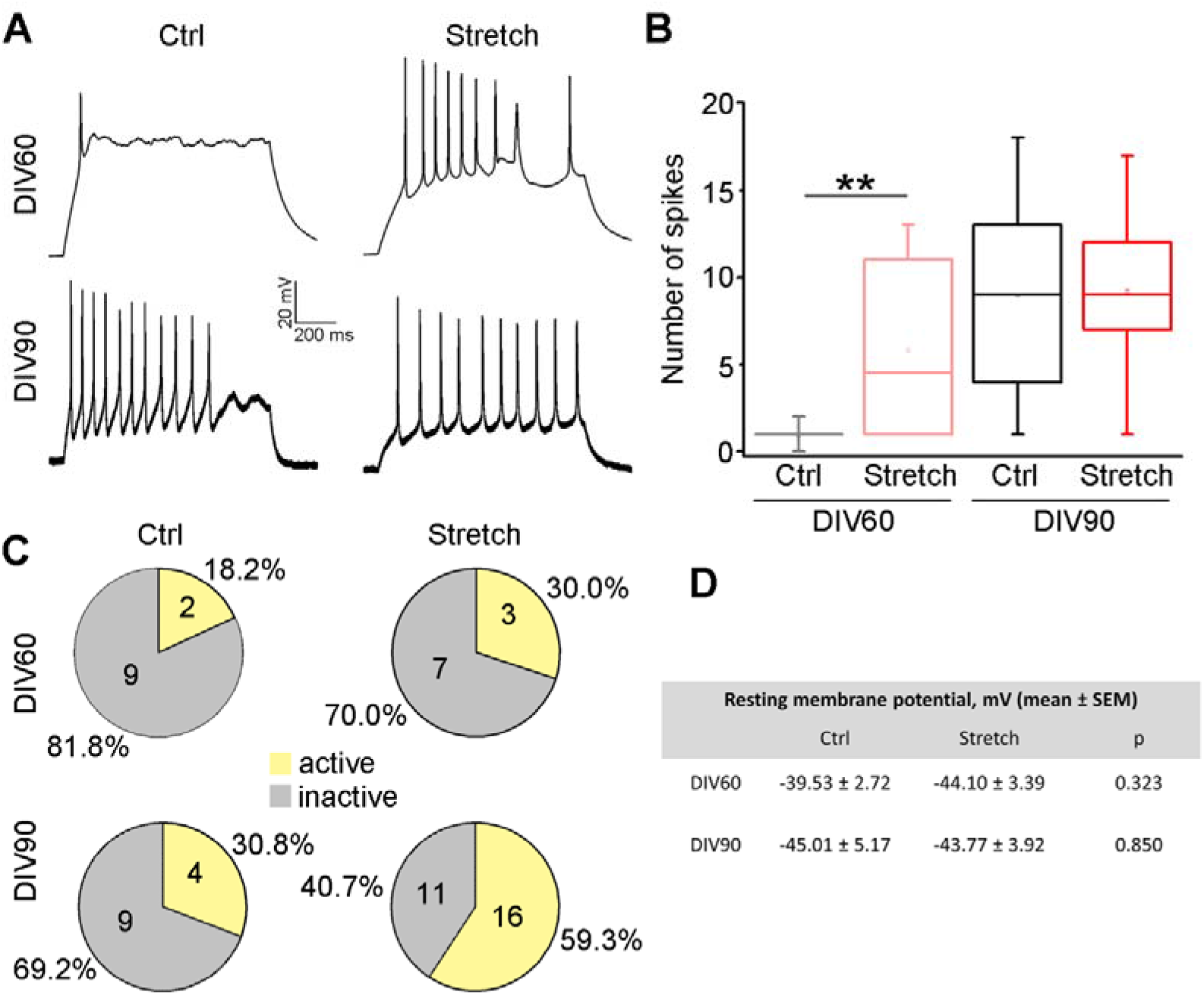
Patch-clamp recordings on SC-NES cell-derived neurons subjected to nano-pulling. A) Representative traces from current clamp recordings of SC-NES cell. B) Quantification of the number of action potentials fired by cells in response to a depolarizing current step (DIV60-ctrl, n=9; DIV60-stretch, n=10; DIV90-ctr, n=11; DIV90-stretch, n=11; Mann-Whitney rank sum test, *p=0.006). C) Histograms showing the percentage of neurons showing spontaneous action potentials; numbers in each sector report the size of each group. D) Table showing no significant difference in the resting membrane potential of cells (DIV60-ctrl, n=11; DIV60-stretch, n=14; DIV90-ctr, n=11; DIV90-stretch, n=11; Student’s t test).

Overall, these results are in agreement with an accelerated path of SC-NES towards a fully functional neuronal phenotype.

### Nano-pulling induces SG of NES cells transplanted into the spinal cord tissue

An organotypic co-culture model was generated by microinjecting DIV10 MNP-loaded and Dil-labelled human SC-NES cells into the ventral horns of day *ex vivo* (DEV) 4 mouse SC slices, according to the protocol shown in Fig. 8A. SC-NES cells were successfully integrated into the SC slice (Fig. 8B, see also the 3D reconstruction in supplementary movie 1 and 2). At DEV10, the fraction of transplanted human cells positive to the apoptotic marker active Caspase 3 was 2.14 ± 2.90% (n=15 slices, on average 500 cells per slice). SC slices were positioned in the millicell insert radially oriented in the ventral-dorsal direction, in order to be aligned to the direction of the force vector (Fig. 8C). At DEV5, the co-cultures were exposed to the magnetic field (stretch) or to a null magnetic field (ctrl) for 48 hours. H-Nestin neural processes sprouting out from the ventral horns of the SC slices were generally oriented in the direction of the dorsoventral axis *(i.e*., the force vector direction) in stretched co-cultures, but not in the unstretched ones (Fig. 8D1).

**Figure 8.**
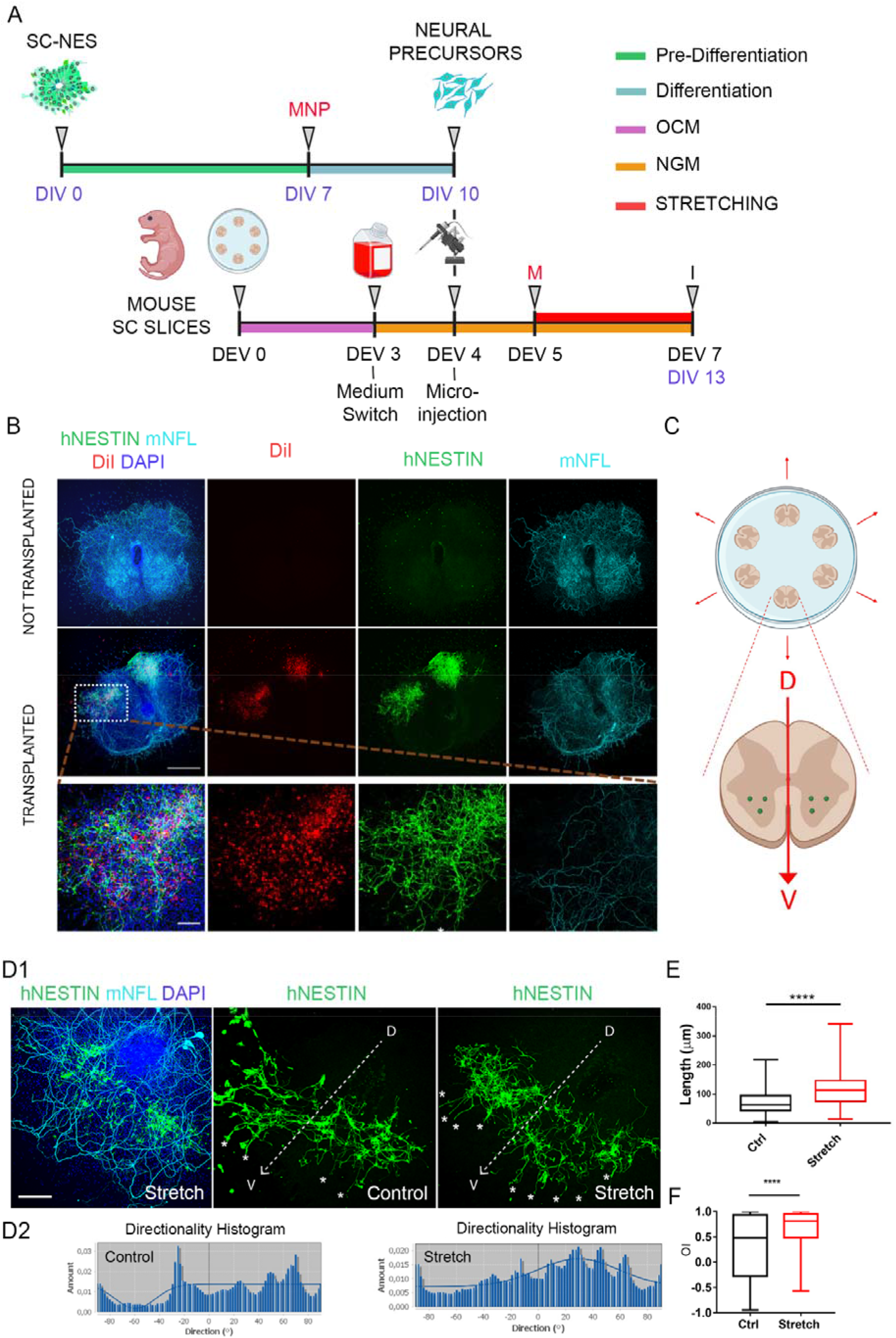
Nano-pulling of human SC-NES cells in mouse SC organotypic slices. (A) Schematic representation of the transplantation protocol of human SC-NES cells loaded with MNPs into mouse SC slices. SC-NES cells are labelled with Dil before transplantation. (B) Representative image of SC slices microinjected with SC-NES cells and non-microinjected (h-Nestin, green; Dil, red; m-NFL, cyan) (DEV10). (C) Schematic representation of co-culture design: up to 6 SC slices are placed concentrically in a millicell with the dorso-ventral axis (red arrow) in the radial direction (centrifugal) and SC-NES cells are microinjected in the ventral horns (3 injections per horn). (D1) SG of SC-NES cells inside the SC slices (h-Nestin, green; m-NFL, cyan; DAPI, blue).) The white arrows show the dorso-ventral axis of the slice. The white stars highlight the presence of processes following a dorso-ventral axis orientation matching the force vector direction. (D2) Directionality histograms for the representative stretched and control co-cultures shown in panel D1. Length (E) and OI (F) of the neural processes sprouting out from the SC slice at DIV 13-14 in stretch and ctrl groups (n=161 processes, n=15 slices, n=4 mice.) Mann Whitney test, p<0.0001. OCM: organotypic culture medium, NGM: neural growth medium (see material & methods).

The direction of h-Nestin neural processes sprouting from the ventral horns of the SC slice was analyzed (Fig. 8D2, relative to the two representative images shown in Fig. 8D1). In stretched co-cultures, neural processes exhibited a preferential orientation, corresponding to the peak of the Gaussian function of the directionality histogram, as documented by the percentage of processes with this preferred orientation (76%) and goodness of fit (0.59). The angle 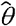 between the preferential orientation and the force vector was < 4°, showing a very high alignment. Conversely, no single preferential orientation for the control co-culture was detected, with a low (i.e., 20%) fraction of processes aligned to the force vector and a goodness of fit of 0.4. A quantitative single-tracing analysis was then performed on the neural processes of h-Nestin positive cells microinjected into SC slices (n>15 slices per group, Fig. 8E). An increase in the length of neural processes in stretch versus control groups was observed (73.16 ± 3.08 μm and 115.40 ± 4.17 μm for ctrl and stretch groups, respectively, p<0.0001, Fig. 8E), resulting in a length increase of approximately 58% after 48 hours of stretching. The distribution of orientation index (Ol=cos *θ* was then calculated, 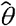 being the angle between the orientation of a neural process and the force vector. The smaller the angle 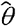 *(i.e*., Ol is approaching to 1), the more aligned to the force vector the process is. A comparison of the data distribution showed a statistically significant increase in the 0l in the stretched versus unstretched condition (p<0.0001).

## DISCUSSION

In this study, by generating active mechanical stimuli, we investigated the involvement of force in the establishment of neuronal cell fate and maturation. Our stretching method is based on cell loading with MNPs and the generation of a dragging force under an external magnetic field.

The net applied force was estimated to be around 10 pN in a cellular process with a length in the order of 100 μm force. With such a small force, a constant load can be applied for hours to months. This makes our method an extraordinary tool for mimicking *in vitro* the condition of continuous loading with the extremely low forces that occur during development.

Short-term stimulation (48 hours) was found to induce a strong elongation which reached the maximum rate of 0.20 ± 0.01 μm·h^−1^·pN^−1^ when NES cells were stimulated between DIV8-10. The elongation took place without thinning (Fig. 2D), indicating that the process is accompanied by the addition of mass.

The same methodology applied to hippocampal neurons was found to induce axon outgrowth at a rate of 0.66 ± 0.02 μm·h^−1^·pN^−1^ (de Vincentiis et al., 2020). The similarity of the response to stretching between mature and immature neurons is not only in terms of elongation. In fact, ultrastructural analyses of immature neural processes of stretched NES cells showed an increase in the linear density of MTs, in the number of tubular ER, and the number of mitochondria. These data are similar findings previously reported for the stretched axons of mouse hippocampal primary neurons (de Vincentiis et al., 2020; Falconieri et al., 2021; Falconieri Alessandro et al., 2022)

This analogy across different stages of neural development raises the question as to how mechanical stimuli represent a well-conserved mechanism to shape the formation of the CNS during development. The observations collected in different models (de Vincentiis et al., 2020; Falconieri Alessandro et al., 2022; V. Raffa et al., 2018) and with different stretching modalities (de Vincentiis et al., 2020; Falconieri et al., 2021) consistently indicate the importance of MTs as a node in signal transduction. MTs of stretched processes are normal in structure but show a statistically significant increase in their density (p<0.0001, Fig. 3A). MTs behave like tension sensors (Hamant et al., 2019), which self-reorganize in the direction of maximal stretch and are stabilized by the application of a tensile force (Putnam et al., 2001). In light of this evidence, we speculate that nano-pulling is perceived by the mechanosensitive MTs, resulting in the stabilization of long stationary MTs (Falconieri Alessandro et al., 2022). A connection between an increase in MT density and mass addition is expected, considering that MTs are the tracks of axonal transport.

During the development of the neural processes of animal cells, peripheral ER usually co-aligns with MTs (Farías et al., 2019; Kimura et al., 2017; Terasaki et al., 1986), showing features such as anterograde and retrograde transport along axonal MTs. As expected, stretched neural processes showed a significant accumulation of tubular ER (p<0.0001, Fig. A4 and B2), similarly to previous observations on hippocampal neurons (de Vincentiis et al., 2020; Falconieri et al., 2021; Falconieri Alessandro et al., 2022). Furthermore MT-based transport of mitochondria into neural processes could explain the observed increase in the number of mitochondria found in the stretched neural processes of human NES cells (Fig. 3A2), as previously reported for mouse primary neurons (Falconieri Alessandro et al., 2022). Our TEM data on NES cells also suggest that, although the mitochondrial morphology is quite similar in the two experimental classes, their length slightly, but significantly, increases upon SG. Together with mitochondria, ER plays a key role in the biosynthesis and calcium buffering in neural processes, which supports axonal transport and mass addition (Markovinovic et al., 2022). Since MT dynamics are also coupled to the axonal transport of vesicles (Yogev et al., 2016), an increase in the total number of available tracks is likely directly correlated with an increase in the total number of transported vesicles. Quantification of vesicles positive to the pre-synaptic marker synaptophysin was performed to support this hypothesis. Pre-synaptic proteins are generally transported in the form of precursor vesicles from the neuronal soma, where they are synthetized, along axonal MTs to the synaptic terminal, where they anchor (Guedes-Dias & Holzbaur, 2019). An increase in the signal was found after 8 and 22 days of stretching (p=0.0003 and p<0.0001, Fig. 6B and 6C, respectively) which reached a plateau after 52 days of stretching (p=0.91, Fig. 6D).

In the light of these observations, we tested the effect of long-term stimulation in depth, by applying a continuous loading for up to 52 days. Long-term stimulation was found to induce remodeling of the cell network (Fig. 4), which was accompanied by functional changes. Specifically, the colocalization between pre- and post-synaptic markers, considered a hallmark of the presence of mature synapses, showed a strong increase after 22 days of stimulation (p<0.0001, Fig. 6A-D). In line with this, TEM imaging revealed an increased percentage in MT-positive cells (p=0.0001, Fig. 6B) in stretched neural processes, indicating later stages of *in vitro* neurogenesis (Haendel et al., 1996).

These results were confirmed by electrophysiological recordings that showed an increase in the number of spikes after 52 days of stretching, reaching a plateau after 90 days of stretching (p=0.006, Fig. 7B). In fact, the expression of synaptic markers and synapse density was high at early stages (DIV30) of maturation, while spiking activity increased at an intermediate stage (DIV60). This difference was lost at a later stage (DIV90) when also the control cultures reached the same maturation level. This mismatch agrees with observations on human stem cell-derived neurons, in which synaptogenesis and structural modifications of the synapse precedes the acquisition of synaptic function (Odawara et al., 2016; Togo et al., 2021; Wilson & Newell-Litwa, 2018). Interestingly, the mechanical stimulation not only speeds up neuronal maturation but also modulates the neuronal fate, as highlighted by the increase in RBFOX3-positive cells after 52 days of stretching (p=0.0008, Fig. 5B3).

Previous studies analyzing the effect of active force application suggested that mechanical passive and active forces may act on the neurodevelopmental trajectories of NSCs by accelerating the maturation of the derived neurons (Arulmoli et al., 2015; Chang et al., 2013; Dai et al., 2019). Here, a chronic stimulation was performed for up to 52 days which clearly demonstrated that mechanical stimulation can approximately halve the time required *in vitro* for the differentiation of neural precursor cells into mature neurons, thus also increasing the number of differentiated cells.

Although the picture is not yet complete, a crucial point arising from current and previous studies (de Vincentiis et al., 2020; Falconieri et al., 2021; Falconieri Alessandro et al., 2022; V. Raffa et al., 2018) is that mechanical stimulation is a well-conserved mechanism that induces a local remodeling in the neural process cytoskeleton. This then triggers the transport and localization of components involved in neural outgrowth, differentiation, and synapse maturation.

As a final phase of this work, experiments were designed to assess the translational potential of the nano-pulling protocol by addressing the complexity of a neural tissue in an *ex vivo* paradigm. We used an organotypic model consisting of SC slices (Falconieri et al., 2021; Pinkernelle et al., 2015; Riggio et al., 2013), onto which we transplanted MNP-loaded human SC-NES cells. After cell engraftment, the co-culture was subjected to nano-pulling, demonstrating that the mechanical stimuli externally applied can induce the guided outgrowth of the neural processes of transplanted NPCs in this complex neural tissue environment. Specifically, the stretched processes showed a strong increase in the elongation rate (2.40 μm·h^−1^ versus 1.52 μm·h^−1^ of the control group) and a strong change in the orientation index, with the stretched processes preferentially aligned in the direction of the force vector (Fig. 8E).

### Ideas and speculations

We speculate that nano-pulling could be explored for the potential application in cell therapies. We are currently testing its use in pre-clinical models of spinal cord injury as a strategy to stimulate SC-NES cell differentiation into mature neurons and thereby to increase the efficiency of their integration into lesioned spinal circuits.

## ACKNOWLEDGMENTS

The study was supported by the Wings for Life Foundation (WFL-IT-16/17 and 20/21), the Italian Ministry of Economic Development through the MAECI (MagNerv), the HFSP (RGP0026/2021). The authors would like to thank Elena Capitanini for re-analysing experimental data and Alessandro Falconieri for the support with XONA chip.

## MATERIAL AND METHODS

### Ethical Statement

All NES cell work was performed according to NIH guidelines for the acquisition and distribution of human tissue for bio-medical research purposes and with approval by the Human Investigation Committees and Institutional Ethics Committees of each institute from which samples were obtained. Final approval from the Committee on Bioethics of the University of Pisa was obtained (Review No. 29/2020). De-identified human specimens were provided by the Joint MRC/Wellcome Trust grant (099175/Z/12/Z), Human Developmental Biology Resource (www.hdbr.org). Appropriate informed consent was obtained, and all available non-identifying information was recorded for each specimen. Tissue was handled in accordance with ethical guidelines and regulations for the research use of human brain tissue set forth by the NIH (http://bioethics.od.nih.gov/humantissue.html) and the WMA Declaration of Helsinki (http://www.wma.net/en/30publications/10policies/b3/index.html).

Animal procedures were performed in strict compliance with protocols approved by Italian Ministry of Public Health and the local Ethical Committee of University of Pisa, in conformity with the Directive 2010/63/EU (project license n°39E1C.N.5Q7 released on 30/10/2021). C57BL/6J mice were kept in a regulated environment (23 ± 1 °C, 50 ± 5% humidity) with a 12 hours light-dark cycle with food and water *ad libitum*.

### Maintenance and differentiation of NES cell lines

Human SC-NES were previously derived as already reported from developing spinal cord tissue(Dell’Anno et al., 2018; Onorati et al., 2016) and NES-iPS cells were derived from human induced pluripotent stem cells (hiPSCs) as already reported (Morelli et al., 2021; Onorati et al., 2016; Sousa et al., 2017). Briefly, these cells hiPSCs were dissociated into single cells in StemFlex medium (#A3349201, Thermo Fisher Scientific, Waltham, Massachusetts)] in Matrigel coated dishes containing 10 μM Y-27632 (#72308, StemCell Technologies, Vancouver, Canada), until confluent. Then, the dual SMAD inhibition protocol was performed, changing the StemFlex medium with a neural induction medium [1:1 Dulbecco’s minimum essential medium/F12 (DMEM/F12) (#11330-032, Gibco, Waltham, Massachusetts)) and Neurobasal medium (#21103-049, Gibco) with addition of B27 supplement (1:50, #175040-44, Gibco), N2 supplement (1:100, #17502-048, Gibco), 20 μg·ml^−1^ insulin (#I9278, Sigma, St. Louis, Missouri), L-glutamine (1:100, #25030-081, Gibco), MEM Non-Essential Amino Acids (1:100, #11140-050, Gibco,) and 2-mercaptoethanol (1:1000, #21985, Gibco)], supplemented with 100 nM of LDN-193189 (# 72144, StemCell Technologies), 10 μM of SB-431542 (#616464-5MG, Merck, Darmstadt, Germany) and 2 μM of XAV939 (#72674, StemCell Technologies). The medium was changed daily until day 11. At day 12, the cells were dissociated with Accutase and maintained in the NES medium [DMEM/F12 with addition of B27 supplement (1:1000), N2 supplement (1:100), 20 ng·ml^−1^ FGF-2 (#13256029, Gibco), 20 ng·ml^−1^ EGF (#PHG0311, Gibco), 1.6 mg·ml^−1^ glucose, 20 μg·ml^−1^ insulin, and 5 ng·ml^−1^ BDNF (#PHC7074, Gibco)] with Y-27632 (10 μM).

Concerning, the NES cells, they were maintained in NES medium [Dulbecco’s minimum essential medium/F12 (DMEM/F12) (#11330-032, Gibco) with addition of B27 supplement (1:1000, #175040-44, Gibco), N2 supplement (1:100, #17502-048, Gibco), 20 ng·ml^−1^ FGF-2 (#13256029, Gibco), 20 ng·ml^−1^ EGF (#PHG0311, Gibco), 1.6 mg-ml^−1^ glucose, 20 μg·ml^−1^ insulin (#I9278, Sigma) and 5 ng·ml^−1^ BDNF (#PHC7074, Gibco)]. Cells were cultured in T25 flasks coated with POLFN [0.01% poly-L-ornithine (#P4957, Sigma), 5 μg·ml^−1^ laminin (#23017-015, Invitrogen, Waltham, Massachusetts) and 1 μg·ml^−1^ fibronectin (#354008, Corning, Corning, New York)] and maintained at 37 °C in a saturated humidity atmosphere of 95% air and 5% CO_2_. Cells were split 1:2 when confluent (~0.5-1 × 10^5^ cells·cm^−2^), once every 4-6 days with 0.25% trypsin. Half volume of the medium was changed every 2-3 days.

Neuronal differentiation of NES cells was performed in two steps. For the pre-differentiation step, cells were seeded in NES medium without FGF-2 and EGF for 7 days, changing the medium every 2-3 days. For the terminal differentiation, cells were dissociated and replated at a density of 0.8-1 × 10^5^ cells·cm^−2^ in a medium composed of DMEM/F12 (1:2), Neurobasal (1:2), N-2 (1:200), B-27 (1:100), Insulin (10 μg·ml^−1^), L-glutamine (1:100) and BDNF (30 ng·ml^−1^). Half volume of the medium was changed every 2-3 days and neurons were differentiated up to 4 months.

### Organotypic model

An organotypic mouse model consisting of spinal cord (SC) slices was used, as previously described (Falconieri et al., 2021; Pinkernelle et al., 2015; Riggio et al., 2013). Spinal cords of P3 mice were dissected under a stereomicroscope in a solution of D-glucose 6.5 mg·ml^−1^ in DPBS (#14190-094, Gibco), sliced (300 μm thickness) and placed on a Millicell membrane (#PICM0RC50, Merck), previously coated with a water solution of 0.1 mg·ml^−1^ collagen (#C7661, Merck), 0.01 mg ml^−1^ poly-L-lysine (#P4707, Merck) and 0.01 mg·ml^−1^ laminin (#L2020, Merck). Then, the SC slices were initially cultured in the organotypic culture medium (OCM) composed of MEM with 25% of horse serum, HBSS and HEPES, 35 nM of D-glucose, 1% penicillin, 1% streptomycin and 2 mM of Glutamax and 0.1 μg·ml^−1^ GDNF (#SRP3200, Merck) (Vyas et al., 2010).

NES cells at early stages of differentiation (DIV10-11) were micro-injected in the ventral horns of the SC slice at DEV 4. Before microinjection, NES cells were stained with 1 μg·ml^−1^ Dil red dye (#C7001, Invitrogen) (5 minutes incubation at 37°C, 15 min incubation at 4°C). After washing with DPBS, cells were detached, counted and resuspended at a concentration of 50 cells per nl in neuronal growth medium (NGM), composed by Neurobasal, N-2 (1:200), B-27 (1:150) and GDNF (100 ng·ml^−1^). 4 nl of cell suspension was microinjected into six locations for each slice (3 injection sites for each ventral horn). Then, SC slices were incubated in NGM and fixed at DEV7.

### MNPs and magnetic field

MNPs used in this study are magnetite MNPs (#4115, Chemicell, Berlin, Germany) with a core of iron oxide ~75 ± 10 nm in diameter and saturation magnetization of 59 Am^2^·kg^−1^, as stated from the supplier. The outer layer is made of glucuronic acid and the hydrodynamic diameter is 100 nm. The intracellular iron was quantified with an iron assay kit (#DIFE-250, BioAssay Systems, Hayward, California), and the absorbance was measured at a wavelength of 590 nm.

Experiments were conducted in 35 mm Petri dishes placed inside a Halbach-like cylinder magnetic applicator, which provided a constant magnetic field gradient of 46.5 T·m^−1^ in the radial centrifugal direction (V. Raffa et al., 2018; Riggio et al., 2014). The force generated by MNPs was calculated by using the formula described in (Riggio et al., 2014):

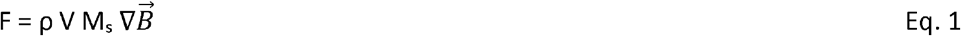

with ρ being the density of iron oxide in the MNP core, V the total volume occupied by the iron core of the MNPs in the neural process, M_s_ the saturation magnetization, and 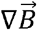 the magnetic field.

### Stretching assay

Two different protocols of stretching were carried out. For the short-term assay cells were dissociated and replated at a density of 1 × 10^5^ cells cm^−2^ in ibidi dish (#80416, IBIDI, Gräfelfing, Germany) precoated with Matrigel (1:60) or POLFN at day *in vitro* (DIV)10, 30, 60, and 90. Alternatively, at DIV10, for confocal and electron imaging, 1-0.5 × 10^5^ cells·cm^−2^ were seeded on a 8 mm glass coverslips (#CB00080RA120MNZ0, Epredia, Portsmouth, New Hampshire) or 2 × 10^5^ cells in microfluidic XONA devices (#RD150, XONA, Research Triangle Park, North Carolina) mounted on 22 glass coverslips (#vntondi.22, F. Franceschini s.a.s., Pisa, Italy) pre-coated with 500 µg·ml^−1^ poly-L-lysine (#P4707, Merck) and 0.01 mg·ml^−1^ laminin (#L2020, Merck). 5 µg·ml^−1^ of MNPs were added 4 hours after seeding. 20 hours later, the Petri dish (stretch group) was put inside the magnetic applicator or kept outside (ctrl group). After 48 hours of stretching, samples were fixed and stained for 15 minutes with 0.1% (vol/vol) crystal violet in ethanol 10% or, alternatively, stained for immunofluorescence or for electron microscopy.

For the long-term assay, cells were seeded for the terminal differentiation at a density of 0.5 × 10^5^ cells·cm^−2^ on 8 mm glass coverslips precoated with POLFN. 5 µg ml^−1^ of MNPs were added 4 hours after seeding and, then, every 15 days up to 4 months. 20 hours after MNP’s addition, the petri dish (stretch group) was put inside the magnetic applicator, keeping outside the control dish (ctrl group). After 9, 23, 53, and 83 days of stretching samples were electrophysiologically recorded or fixed and stained for immunofluorescence.

For the organotypic model, 4-6 slices were placed in a millicells with the dorsal-ventral axis in the radial direction (outward orientation) (Fig. 8C)

### Immunostaining

For immunostaining, NES cells were washed with PBS and fixed with 4% formaldehyde (FA) for 12 minutes at room temperature (RT). After three washes of 3 minutes with PBS-Triton X-100 (PBSX) [0.1% (vol/vol) Triton X-100 in PBS Ca^2+^/Mg^2+^ 1X], samples were permeabilized with permeabilization solution [0.5% (vol/vol) Triton X-100 in PBS Ca^2+^/Mg^2+^ 1X] and blocked at RT for 1 hour with blocking solution [5% fetal bovine serum, 0.3% (vol/vol) Triton X-100 in PBS Ca^2+^/Mg^2+^ 1X]. Primary antibodies were diluted in antibody solution [3% fetal bovine serum, 0.2% (vol/vol) Triton X-100 in PBS Ca^2+^/Mg^2+^ 1X] and incubated overnight at 4 °C. Primary antibodies were diluted as follows: TUBB3 (1:500, #T8578, Sigma), Synaptophysin (1:500, #Mab329, Millipore, Burlington, Massachusetts), Homer 1b/c (1:350, #160023, Synaptic System, Göttingen, Germany), KDEL (1:200, PA1-013, Invitrogen,), SOX2 (1:400, ab5603, Millipore,) RBFOX3 (1:500, #ABN78, Millipore), NFL (1:200, #2424662, Millipore), and Nestin (1:200, MAB1259, R&D, Minneapolis, Minnesota). The next day, samples were washed three times for 3 minutes and then incubated 1 hour at RT with secondary antibody (1:500, #A21202, #A21449, #A10042, #A32731, Life Technologies, Carlsbad, California) and Hoechst 33342 (1:1000, #H3570, Invitrogen,) or DAPI (1:1000, #32670, Sigma). After two washes of 3 minutes with PBSX and one wash of 3 minutes with PBS Ca^2+^/Mg^2+^, the fixed cells were directly imaged or mounted on microscope slides.

SC slices were fixed in 4% FA for 30 min at RT, washed three times in PBS and permeabilized (0.7% Triton in PBS) for 10 min at RT. Blocking (Triton 0.5%+FBS 10% in PBS) was performed for 4h at 4°C. Membranes were incubated overnight at 4°C in primary antibody solution (Triton 0.5%+FBS 1% in PBS + primary antibody: neurofilament (1:500, #840801, COVANCE, Princeton, New Jersey), Nestin (1:200, #MAB1259, R&D,). Then, the samples were washed three times in PBS and incubated in secondary antibody solution (Triton 0.5%+FBS 1% in PBS + secondary antibody at working dilution 1:500) for 3h at RT in dark condition. Slices were washed three times in PBS for 10 min, mounted on glass slices by using Aqua-Poly/Mount solution and examined using a confocal microscope.

All images were acquired using a fluorescent microscope (TE2000-U, Nikon, Tokyo, Japan) equipped with DS-Ri2 camera or a laser scanning confocal microscope (Eclipse Ti, Nikon). With the fluorescent microscope, images were acquired with a 10x objective and 1024 × 1024 pixel resolution, while 60x and 20x objective oil immersion were used with the laser scanning confocal microscope. Here, series of ◻ 40 optical plans in Z were acquired at 1024 × 1024 pixel resolution with a z-step of 0.2 μm. Images were acquired with a 405 nm laser (425-475 emission filter) or a 488 nm laser (500-550 emission filter) or a 561 laser (570-620 emission filter) or a 640 nm laser (663-738 emission filter) and using an exposure time of 100 ms.

### Transmission electron microscopy (TEM) analysis

NES cells, plated on 8 mm glass coverslips at the density of 1 × 10^5^ cells·cm^−2^, were treated as previously described (de Vincentiis et al., 2020). Briefly, cells were fixed with 1.5% glutaraldehyde in Na Cacodylate buffer (0.1 M, pH 7.4), washed in the same buffer and postfixed with reduced osmium tetroxide solution (1% K_3_Fe(CN)_6_ + 1% OsO_4_ in Na Cacodylate buffer). After rinses, NES cells were stained with × solution diluted 1:10 (v/v) in 20% ethanol/water (Moscardini et al., 2020), then dehydrated with a series of ethanol solutions of increasing concentration. Cells were finally embedded in epoxy resin (Epoxy embedding medium kit, Merck KGaA) that was then baked for 24 hours at 60 °C. After parting resin from coverslips, samples were sectioned with UC7 ultramicrotome (Leica Microsystems, Wetzlar, Germany) in 80 nm thicker sections and collected on 300 mesh copper grids (G300Cu - Electron Microscopy Science). Grids were finally analyzed with a Zeiss Libra 120 Plus transmission electron microscope, operating at 120 kV and equipped with an in-column omega filter (for the energy filtered imaging) and 16-bit CCD camera 2 k × 2 k bottom mounted (Zeiss, Oberkichen, Germany). 8000X magnification micrographs were collected for the quantitative analysis of cells ultrastructure.

### Electrophysiology

SC-NES-derived neurons were recorded by adapting the protocols described in (de Vincentiis et al., 2020). Cultures were continuously perfused with oxygenated Tyrode’s solution containing (in mM): NaCl 150, KCl 4, MgCl_2_ 1, CaCl_2_ 4, Glucose 10, HEPES 10, pH 7.4 with NaOH. Borosilicate glass pipettes were pulled to a resistance of 4-6 MΩ using a PC-100 puller (Narishige, Japan) and filled with an internal solution containing (in mM): K-Gluconate 145, MgCl2 2, HEPES 10, EGTA 0.1, Mg-ATP 2.5, Na-GTP 0.25, phosphocreatine 5, pH 7.35 with KOH. After achieving a stable whole-cell configuration, the amplifier was switched to current clamp mode and the holding current was adjusted to have an initial membrane potential of −70 mV. Then, a depolarizing step of 20-25 pA was applied and the number of action potentials fired was recorded. The resting membrane potential and spontaneous spiking activity were measured for at least 1 min in I=0 configuration. Data were acquired using a MultiClamp 700A amplifier, connected to a Digidata 1550B digitizer (Molecular Devices, San Jose, California) and analyzed using Clampfit 11.2 (Molecular Devices).

### Image analysis

In short-term stretching assay, the analyses were performed by tracing single neural process. The elongation was measured by using the plugin NeuronJ (Popko et al., 2009). 200 non-interconnected TUBB3-stained neural processes (cut-off 20 µm) were analyzed from 10× magnification images (randomly acquired). For process thickness, a population of 40 crystal violet-stained neural processes was analyzed from randomly acquired 10× magnification images. For each cell, the longest neural process l has been considered and measured. After threshold normalization, binary conversion, the occupied area A related to that neural process has been estimated and the thickness *s* was calculated as *s* = *A/l*. For the analysis of fluorescence, from 60× magnification images (randomly acquired), the integrated density of each cell was measured with the image analysis software ImageJ, and normalized for the relative area calculated after threshold normalization.

For long-term stretching assay, network analysis methods were chosen. The area of the neural process network was evaluated from 10× magnification images (randomly acquired) obtained from a composition of four 20× images. For each image, the network area was calculated as the ratio of the area occupied by neural processes (after nuclei’s area subtraction) and the number of nuclei, automatically counted with the function “analyze particle” of Fiji software. For the analysis of cell dispersion, the center of mass of all nuclei was measured from 10× randomly acquired magnification images obtained from a composition of four 20× images. For each image, the presence of the nuclei was detected thanks to the “Analyze particles” function of ImageJ software, after threshold setting. The center of mass of all nuclei was measured. Subsequently, for each nucleus, the distance between its center of mass and the one of the nearest nucleus was calculated and the dispersion analyzed. In order to provide an evidence of different network morphology, the mean node distance 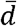 was defined as:

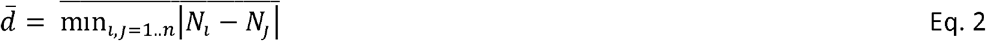

With a network of nodes *N* (center of mass of each cell soma) and arcs *d* (distance between nodes), being *n* the total number of nodes.

The fluorescence of the neural process network was evaluated from 60× magnification images randomly acquired. The fluorescence as the integrated density of the neural processes was measured with ImageJ software and normalized for the relative network’s area, after nuclei subtraction. The synapse density was estimated by evaluating the number of colocalization spots of pre- and post-synaptic markers, individuated by the plugin puncta analyzer, as previously described (Ippolito & Eroglu, 2010). Briefly, both channels were converted using the maximum intensity of the Z projection and merged. Following the identification of a specific ROI corresponding to a process of the cell, and keeping default parameters, the plugin quantifies the puncta in each channel and the co-localized puncta between the two channels. The obtained information was then normalized for the considered area. 63× magnification images were used to perform this analysis. Cellular differentiation was determined by manually counting individual cells labeled for Sox2 or RBFOX3 and expressing them as a percentage of the total number of cells in the field (*e.g.*, positive for DAPI). To perform this analysis, 10× magnification images (randomly acquired) obtained from a composition of four 20× images were used.

### TEM analysis

TEM analysis was performed using ImageJ and the plugin NeuronJ (Meijering et al., 2004). Particularly, for the evaluation of ER) density, the length of ER cisternae recognized as electro-dense and oblong structures was traced and measured with NeuronJ, and normalized for the total area of the neural process considered. For microtubule density quantification, the number of microtubules identified as tubular structures was manually counted in an organelle-free region of the neural process. Then, the obtained value was normalized for the diameter of the corresponding region of the neural process, giving one value per neural process. For the cytoskeletal composition the same data were used, comparing the number of neural processes with microtubules and the ones without (*e.g.* with microfilaments).

### Statistical analysis

Data are reported as mean ± standard deviation (SD) or standard error of the mean (SEM) from at least four separate experiments after blinded analyses (except immunostaining against SOX2/RBFOX3: three separate experiments; TEM data: two separate experiments). Data were plotted with GraphPad software, version 6.0. The normality of the distribution was assayed by different tests, such as D’Agostino & Pearson normality test, Shapiro-Wilk normality test, or KS normality test. For normally distributed data, one-way, two-way analysis of variance (ANOVA) test or *t*-test for unpaired data followed by Bonferroni correction were used. For non-normally distributed data, Kolmogorov-Smirnov test, Mann Whitney, or Kruskal-Wallis test analyses were carried out. Significance was set at *p* ≤ 0.05.

### Data availability

All metadata, data and data analysis are available at: https://drive.google.com/drive/folders/1xROQdtX8bbxPmAJ3LStyB1wVYyiyKC8N?usp=sharing. Our team adhere to the FAIR criteria on Findability, Accessibility, Interoperability, and Reuse of scientific data.

---

**Figure S1.**
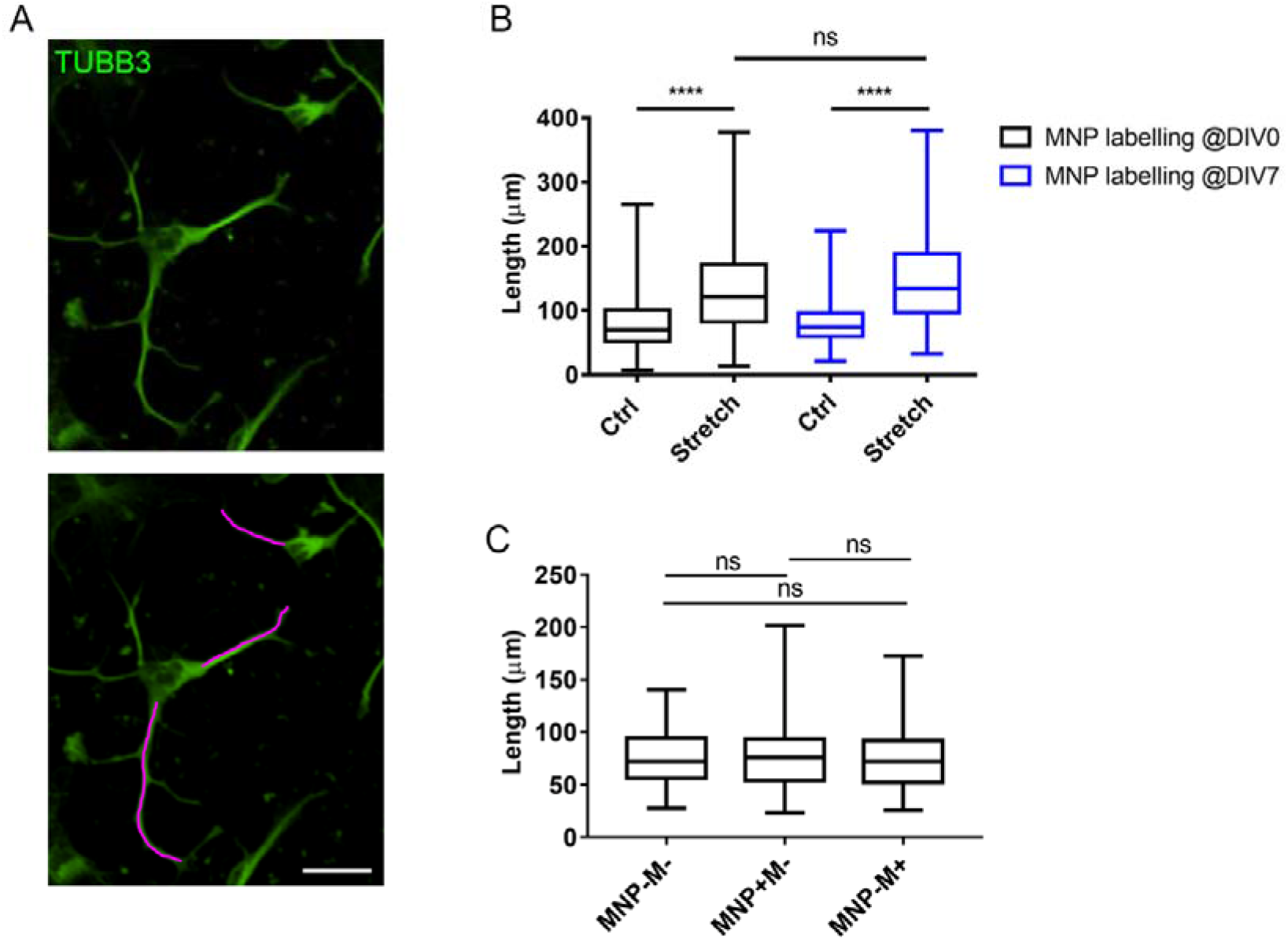
**(A)** Representative images of SC-NES cells for process elongation analysis. Primary processes are traced (magenta) and measured with the NeuonJ plugin. Scale bar: 20 µm. **(B)** Process length of SC-NES cells at DIV10. Comparison between control and stretched samples with MNP labelling @DIV0 (black) and control and stretched samples with MNP labelling @DIV7 (blue). Box plot, min-to-max, n=400 neural processes. Kruskal-Wallis test with Dunn post hoc test, p<0.0001. **(C)** Process length of SC-NES cells at DIV10. Comparison between control conditions. Box plot, min-to-max, n=100. Kruskal-Wallis test with Dunn post hoc test, p=0.8613.

**Figure S2.**
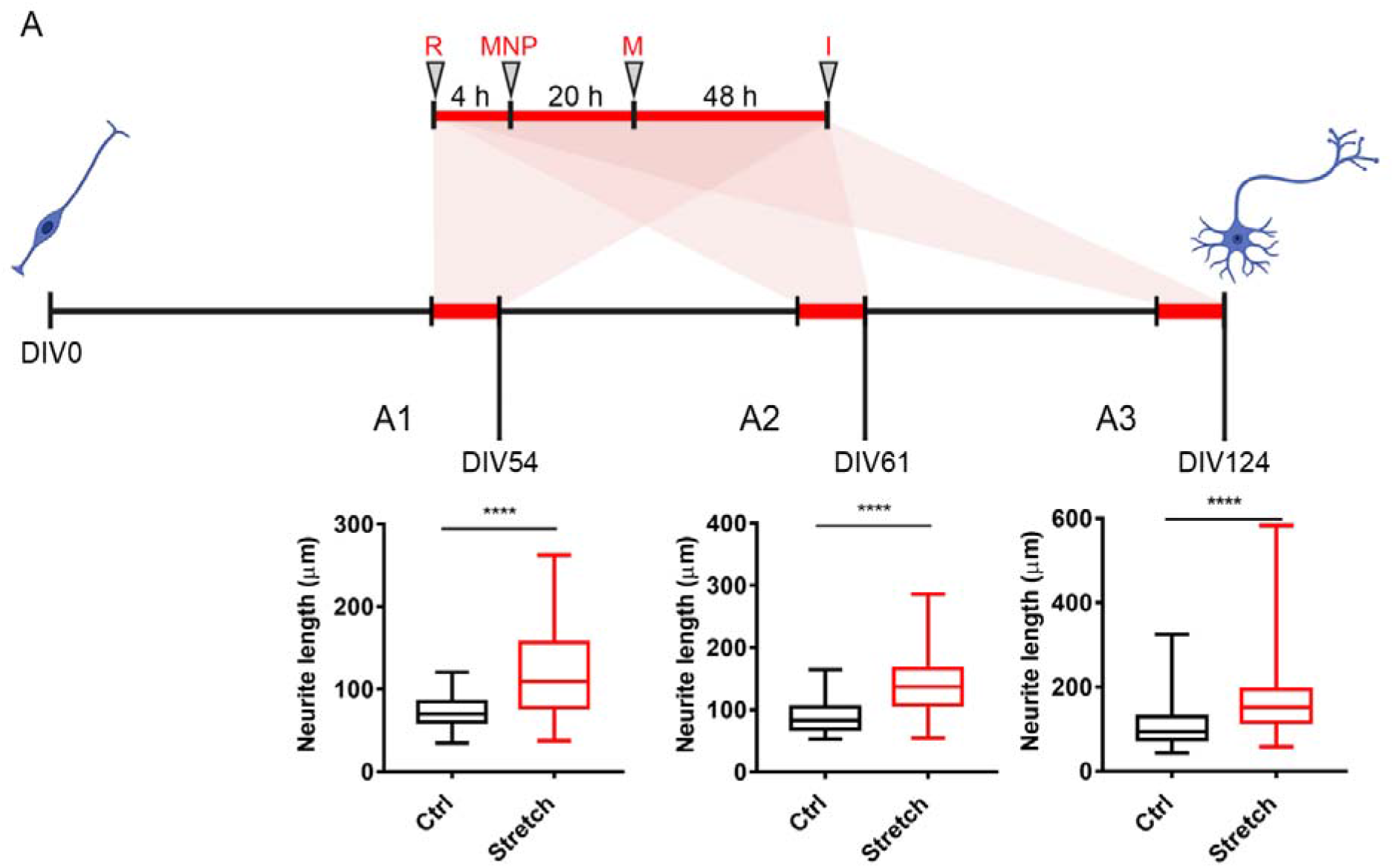
Short-term stretching assay of NES-iPS cells. (A) Schematic representation of the experimental design. At DIV51, 58 and 121 cells are replated (R), 4 h later MNPs are added, 20 h later the “stretch” groups are placed inside a magnetic applicator (M). After 48 h of continuous stretching samples are fixed and subjected to imaging (I). (A1) Neural process length at DIV54. Box plot, min-to-max, n=50. T-test for unpaired data, p<0.0001, t=5.597, df=98; (A2) Neural process length at DIV61. Box plot, min-to-max, n=50. Mann-Whitney test, p<0.0001. (A3) Neural process length at DIV124. Box plot, min-to-max, n=200. Mann-Whitney test, p<0.0001.

Supplementary movie 1-2: integration of transplanted SC-NES in the SC slice.

## BIBLIOGRAPHY

Abdel Fattah, A. R., & Ranga, A. (2020). Nanoparticles as Versatile Tools for Mechanotransduction in Tissues and Organoids. Frontiers in Bioengineering and Biotechnology, 8. https://doi.org/10.3389/fbioe.2020.00240

Abuwarda, H., & Pathak, M. M. (2020). Mechanobiology of neural development. Current Opinion in Cell Biology, 66, 104–111. https://doi.org/10.1016/j.ceb.2020.05.012

Arulmoli, J., Pathak, M. M., McDonnell, L. P., Nourse, J. L., Tombola, F., Earthman, J. C., & Flanagan, L. A. (2015). Static stretch affects neural stem cell differentiation in an extracellular matrix-dependent manner. Scientific Reports, 5(1), 8499. https://doi.org/10.1038/srep08499

Barkho, B. Z., Song, H., Aimone, J. B., Smrt, R. D., Kuwabara, T., Nakashima, K., Gage, F. H., & Zhao, X. (2006). Identification of Astrocyte-expressed Factors That Modulate Neural Stem/Progenitor Cell Differentiation. Stem Cells and Development, 15(3), 407–421. https://doi.org/10.1089/scd.2006.15.407

Breunig, J. J., Haydar, T. F., & Rakic, P. (2011). Neural Stem Cells: Historical Perspective and Future Prospects. Neuron, 70(4), 614–625. https://doi.org/10.1016/j.neuron.2011.05.005

Chang, Y.-J., Tsai, C.-J., Tseng, F.-G., Chen, T.-J., & Wang, T.-W. (2013). Micropatterned stretching system for the investigation of mechanical tension on neural stem cells behavior. Nanomedicine: Nanotechnology, Biology and Medicine, 9(3), 345–355. https://doi.org/10.1016/j.nano.2012.07.008

Chatelin, S., Constantinesco, A., & Willinger, R. (2010). Fifty years of brain tissue mechanical testing: From in vitro to in vivo investigations. Biorheology, 47(5–6), 255–276. https://doi.org/10.3233/BIR-2010-0576

Chklovskii, D. B. (2004). Synaptic Connectivity and Neuronal Morphology. Neuron, 43(5), 609–617. https://doi.org/10.1016/j.neuron.2004.08.012

Dai, R., Hang, Y., Liu, Q., Zhang, S., Wang, L., Pan, Y., & Chen, H. (2019). Improved neural differentiation of stem cells mediated by magnetic nanoparticle-based biophysical stimulation. Journal of Materials Chemistry B, 7(26), 4161–4168. https://doi.org/10.1039/C9TB00678H

de Vincentiis, S., Falconieri, A., Mainardi, M., Cappello, V., Scribano, V., Bizzarri, R., Storti, B., Dente, L., Costa, M., & Raffa, V. (2020). Extremely Low Forces Induce Extreme Axon Growth. Journal of Neuroscience. https://doi.org/10.1523/JNEUROSCI.3075-19.2020

De Vincentiis, S., Falconieri, A., Mickoleit, F., Cappello, V., Schüler, D., & Raffa, V. (2021). Induction of Axonal Outgrowth in Mouse Hippocampal Neurons via Bacterial Magnetosomes. International Journal of Molecular Sciences, 22(8), 4126. https://doi.org/10.3390/ijms22084126

Dell’Anno, M. T., Wang, X., Onorati, M., Li, M., Talpo, F., Sekine, Y., Ma, S., Liu, F., Cafferty, W. B. J., Sestan, N., & Strittmatter, S. M. (2018). Human neuroepithelial stem cell regional specificity enables spinal cord repair through a relay circuit. Nature Communications. https://doi.org/10.1038/s41467-018-05844-8

Discher, D. E., Janmey, P., & Wang, Y. (2005). Tissue Cells Feel and Respond to the Stiffness of Their Substrate. Science, 310(5751), 1139–1143. https://doi.org/10.1126/science.1116995

Essen, D. C. Van. (1997). A tension-based theory of morphogenesis and compact wiring in the central nervous system. Nature, 385(6614), 313–318. https://doi.org/10.1038/385313a0

Falconieri, A., Taparia, N., De Vincentiis, S., Cappello, V., Sniadecki, N. J., & Raffa, V. (2021). MAGNETICALLY-ACTUATED MICROPOSTS STIMULATE AXON GROWTH. Biophysical Journal. https://doi.org/10.1016/j.bpj.2021.12.041

Falconieri Alessandro, Vincentiis Sara De, Cappello Valentina, Convertino Domenica, Ghignoli Samuele, Figoli Sofia, Luin Stefano, Catala-Castro Frederic, Marchetti Laura, Borello Ugo, Krieg Michael, & Raffa Vittoria. (2022). Axonal plasticity in response to active forces generated through magnetic nano-pulling. BioRxiv.

Farías, G. G., Fréal, A., Tortosa, E., Stucchi, R., Pan, X., Portegies, S., Will, L., Altelaar, M., & Hoogenraad, C. C. (2019). Feedback-Driven Mechanisms between Microtubules and the Endoplasmic Reticulum Instruct Neuronal Polarity. Neuron, 102(1), 184–201.e8. https://doi.org/10.1016/j.neuron.2019.01.030

Franze, K. (2013). The mechanical control of nervous system development. Development, 140(15), 3069–3077. https://doi.org/10.1242/dev.079145

Guedes-Dias, P., & Holzbaur, E. L. F. (2019). Axonal transport: Driving synaptic function. Science, 366(6462). https://doi.org/10.1126/science.aaw9997

Haendel, M. A., Bollinger, K. E., & Baas, P. W. (1996). Cytoskeletal changes during neurogenesis in cultures of avian neural crest cells. Journal of Neurocytology, 25(1), 289–301. https://doi.org/10.1007/BF02284803

Hamant, O., Inoue, D., Bouchez, D., Dumais, J., & Mjolsness, E. (2019). Are microtubules tension sensors? Nature Communications, 10(1), 2360. https://doi.org/10.1038/s41467-019-10207-y

Ippolito, D. M., & Eroglu, C. (2010). Quantifying Synapses: an Immunocytochemistry-based Assay to Quantify Synapse Number. Journal of Visualized Experiments, 45. https://doi.org/10.3791/2270

Iwata, R., & Vanderhaeghen, P. (2021). Regulatory roles of mitochondria and metabolism in neurogenesis. Current Opinion in Neurobiology, 69, 231–240. https://doi.org/10.1016/j.conb.2021.05.003

Kalyani, A. J., Piper, D., Mujtaba, T., Lucero, M. T., & Rao, M. S. (1998). Spinal Cord Neuronal Precursors Generate Multiple Neuronal Phenotypes in Culture. The Journal of Neuroscience, 18(19), 7856–7868. https://doi.org/10.1523/JNEUROSCI.18-19-07856.1998

Khacho, M., Harris, R., & Slack, R. S. (2019). Mitochondria as central regulators of neural stem cell fate and cognitive function. Nature Reviews Neuroscience, 20(1), 34–48. https://doi.org/10.1038/s41583-018-0091-3

Kimura, K., Mamane, A., Sasaki, T., Sato, K., Takagi, J., Niwayama, R., Hufnagel, L., Shimamoto, Y., Joanny, J.-F., Uchida, S., & Kimura, A. (2017). Endoplasmic-reticulum-mediated microtubule alignment governs cytoplasmic streaming. Nature Cell Biology, 19(4), 399–406. https://doi.org/10.1038/ncb3490

Kumar, A., Placone, J. K., & Engler, A. J. (2017). Understanding the extracellular forces that determine cell fate and maintenance. Development, 144(23), 4261–4270. https://doi.org/10.1242/dev.158469

Kunze, A., Tseng, P., Godzich, C., Murray, C., Caputo, A., Schweizer, F. E., & Di Carlo, D. (2015). Engineering cortical neuron polarity with nanomagnets on a chip. ACS Nano. https://doi.org/10.1021/nn505330w

Luo, L., & O’Leary, D. D. M. (2005). Axon retraction and degeneration in development and disease. In Annual Review of Neuroscience. https://doi.org/10.1146/annurev.neuro.28.061604.135632

Mandal, S., Lindgren, A. G., Srivastava, A. S., Clark, A. T., & Banerjee, U. (2011). Mitochondrial Function Controls Proliferation and Early Differentiation Potential of Embryonic Stem Cells. Stem Cells, 29(3), 486–495. https://doi.org/10.1002/stem.590

Markovinovic, A., Greig, J., Martín-Guerrero, S. M., Salam, S., & Paillusson, S. (2022). Endoplasmic reticulum–mitochondria signaling in neurons and neurodegenerative diseases. Journal of Cell Science, 135(3). https://doi.org/10.1242/jcs.248534

Meijering, E., Jacob, M., Sarria, J.-C. F., Steiner, P., Hirling, H., & Unser, M. (2004). Design and validation of a tool for neurite tracing and analysis in fluorescence microscopy images. Cytometry, 58A(2), 167–176. https://doi.org/10.1002/cyto.a.20022

Morelli, E., Speranza, E. A., Pellegrino, E., Beznoussenko, G. V., Carminati, F., Garré, M., Mironov, A. A., Onorati, M., & Vaccari, T. (2021). Activity of the SNARE Protein SNAP29 at the Endoplasmic Reticulum and Golgi Apparatus. Frontiers in Cell and Developmental Biology, 9. https://doi.org/10.3389/fcell.2021.637565

Moscardini, A., Di Pietro, S., Signore, G., Parlanti, P., Santi, M., Gemmi, M., & Cappello, V. (2020). Uranium-free X solution: a new generation contrast agent for biological samples ultrastructure. Scientific Reports, 10(1), 11540. https://doi.org/10.1038/s41598-020-68405-4

Nishimura, T., Honda, H., & Takeichi, M. (2012). Planar Cell Polarity Links Axes of Spatial Dynamics in Neural-Tube Closure. Cell, 149(5), 1084–1097. https://doi.org/10.1016/j.cell.2012.04.021

Odawara, A., Katoh, H., Matsuda, N., & Suzuki, I. (2016). Physiological maturation and drug responses of human induced pluripotent stem cell-derived cortical neuronal networks in long-term culture. Scientific Reports, 6(1), 26181. https://doi.org/10.1038/srep26181

Onorati, M., Li, Z., Liu, F., Sousa, A. M. M., Nakagawa, N., Li, M., Dell’Anno, M. T., Gulden, F. O., Pochareddy, S., Tebbenkamp, A. T. N., Han, W., Pletikos, M., Gao, T., Zhu, Y., Bichsel, C., Varela, L., Szigeti-Buck, K., Lisgo, S., Zhang, Y., … Sestan, N. (2016). Zika Virus Disrupts Phospho-TBK1 Localization and Mitosis in Human Neuroepithelial Stem Cells and Radial Glia. Cell Reports. https://doi.org/10.1016/j.celrep.2016.08.038

Pascual-Leone, A., Amedi, A., Fregni, F., & Merabet, L. B. (2005). THE PLASTIC HUMAN BRAIN CORTEX. Annual Review of Neuroscience, 28(1), 377–401. https://doi.org/10.1146/annurev.neuro.27.070203.144216

Pathak, M. M., Nourse, J. L., Tran, T., Hwe, J., Arulmoli, J., Le, D. T. T., Bernardis, E., Flanagan, L. A., & Tombola, F. (2014). Stretch-activated ion channel Piezo1 directs lineage choice in human neural stem cells. Proceedings of the National Academy of Sciences, 111(45), 16148–16153. https://doi.org/10.1073/pnas.1409802111

Pfister, B. J., Iwata, A., Meaney, D. F., & Smith, D. H. (2004). Extreme stretch growth of integrated axons. Journal of Neuroscience. https://doi.org/10.1523/JNEUROSCI.1974-04.2004

Pinkernelle, J., Raffa, V., Calatayud, M. P., Goya, G. F., Riggio, C., & Keilhoff, G. (2015). Growth factor choice is critical for successful functionalization of nanoparticles. Frontiers in Neuroscience, 9(SEP). https://doi.org/10.3389/fnins.2015.00305

Piper, D. R., Mujtaba, T., Keyoung, H., Roy, N. S., Goldman, S. A., Rao, M. S., & Lucero, M. T. (2001). Identification and characterization of neuronal precursors and their progeny from human fetal tissue. Journal of Neuroscience Research, 66(3), 356–368. https://doi.org/10.1002/jnr.1228

Pita-Thomas, W., Steketee, M. B., Moysidis, S. N., Thakor, K., Hampton, B., & Goldberg, J. L. (2015). Promoting filopodial elongation in neurons by membrane-bound magnetic nanoparticles. Nanomedicine: Nanotechnology, Biology and Medicine, 11(3), 559–567. https://doi.org/10.1016/j.nano.2014.11.011

Popko, J., Fernandes, A., Brites, D., & Lanier, L. M. (2009). Automated analysis of NeuronJ tracing data. Cytometry Part A, 75A(4), 371–376. https://doi.org/10.1002/cyto.a.20660

Putnam, A. J., Schultz, K., & Mooney, D. J. (2001). Control of microtubule assembly by extracellular matrix and externally applied strain. American Journal of Physiology-Cell Physiology, 280(3), C556–C564. https://doi.org/10.1152/ajpcell.2001.280.3.C556

Raffa, V., Falcone, F., Calatayud, M. P., Goya, G. F., & Cuschieri, A. (2018). Title: Pico-Newton mechanical forces promote neurite growth. In arXiv.

Raffa, Vittoria. (2022). Force: A messenger of axon outgrowth. Seminars in Cell & Developmental Biology. https://doi.org/10.1016/j.semcdb.2022.07.004

Raffa, Vittoria, Falcone, F., De Vincentiis, S., Falconieri, A., Calatayud, M. P., Goya, G. F., & Cuschieri, A. (2018). Piconewton Mechanical Forces Promote Neurite Growth. Biophysical Journal, 115(10), 2026–2033. https://doi.org/10.1016/j.bpj.2018.10.009

Rammensee, S., Kang, M. S., Georgiou, K., Kumar, S., & Schaffer, D. V. (2017). Dynamics of Mechanosensitive Neural Stem Cell Differentiation. Stem Cells, 35(2), 497–506. https://doi.org/10.1002/stem.2489

Riggio, C., Calatayud, M. P., Giannaccini, M., Sanz, B., Torres, T. E., Fernández-Pacheco, R., Ripoli, A., Ibarra, M. R., Dente, L., Cuschieri, A., Goya, G. F., & Raffa, V. (2014). The orientation of the neuronal growth process can be directed via magnetic nanoparticles under an applied magnetic field. Nanomedicine: Nanotechnology, Biology, and Medicine, 10(7). https://doi.org/10.1016/j.nano.2013.12.008

Riggio, C., Nocentini, S., Catalayud, M. P., Goya, G. F., Cuschieri, A., Raffa, V., & del Río, J. A. (2013). Generation of magnetized olfactory ensheathing cells for regenerative studies in the central and peripheral nervous tissue. International Journal of Molecular Sciences, 14(6). https://doi.org/10.3390/ijms140610852

Shen, Q., Goderie, S. K., Jin, L., Karanth, N., Sun, Y., Abramova, N., Vincent, P., Pumiglia, K., & Temple, S. (2004). Endothelial Cells Stimulate Self-Renewal and Expand Neurogenesis of Neural Stem Cells. Science, 304(5675), 1338–1340. https://doi.org/10.1126/science.1095505

Silbereis, J. C., Pochareddy, S., Zhu, Y., Li, M., & Sestan, N. (2016). The Cellular and Molecular Landscapes of the Developing Human Central Nervous System. Neuron, 89(2), 248–268. https://doi.org/10.1016/J.NEURON.2015.12.008

Sousa, A. M. M., Zhu, Y., Raghanti, M. A., Kitchen, R. R., Onorati, M., Tebbenkamp, A. T. N., Stutz, B., Meyer, K. A., Li, M., Kawasawa, Y. I., Liu, F., Perez, R. G., Mele, M., Carvalho, T., Skarica, M., Gulden, F. O., Pletikos, M., Shibata, A., Stephenson, A. R., … Sestan, N. (2017). Molecular and cellular reorganization of neural circuits in the human lineage. Science, 358(6366), 1027–1032. https://doi.org/10.1126/science.aan3456

Steketee, M. B., Moysidis, S. N., Jin, X. L., Weinstein, J. E., Pita-Thomas, W., Raju, H. B., Iqbal, S., & Goldberg, J. L. (2011). Nanoparticle-mediated signaling endosome localization regulates growth cone motility and neurite growth. Proceedings of the National Academy of Sciences of the United States of America. https://doi.org/10.1073/pnas.1019624108

Stiles, J., & Jernigan, T. L. (2010). The Basics of Brain Development. Neuropsychology Review, 20(4), 327–348. https://doi.org/10.1007/s11065-010-9148-4

Suzuki, M., Morita, H., & Ueno, N. (2012). Molecular mechanisms of cell shape changes that contribute to vertebrate neural tube closure. Development, Growth & Differentiation, 54(3), 266–276. https://doi.org/10.1111/j.1440-169X.2012.01346.x

Temple, S. (2001). The development of neural stem cells. Nature, 414(6859), 112–117. https://doi.org/10.1038/35102174

Terasaki, M., Chen, L. B., & Fujiwara, K. (1986). Microtubules and the endoplasmic reticulum are highly interdependent structures. Journal of Cell Biology, 103(4), 1557–1568. https://doi.org/10.1083/jcb.103.4.1557

Togo, K., Fukusumi, H., Shofuda, T., Ohnishi, H., Yamazaki, H., Hayashi, M. K., Kawasaki, N., Takei, N., Nakazawa, T., Saito, Y., Baba, K., Hashimoto, H., Sekino, Y., Shirao, T., Mochizuki, H., & Kanemura, Y. (2021). Postsynaptic structure formation of human iPS cell-derived neurons takes longer than presynaptic formation during neural differentiation in vitro. Molecular Brain, 14(1), 149. https://doi.org/10.1186/s13041-021-00851-1

Valenzuela, J. I., & Perez, F. (2015). Diversifying the secretory routes in neurons. Frontiers in Neuroscience, 9. https://doi.org/10.3389/fnins.2015.00358

Verstraelen, P., Garcia-Diaz Barriga, G., Verschuuren, M., Asselbergh, B., Nuydens, R., Larsen, P. H., Timmermans, J.-P., & De Vos, W. H. (2020). Systematic Quantification of Synapses in Primary Neuronal Culture. IScience, 23(9), 101542. https://doi.org/10.1016/j.isci.2020.101542

Verstraelen, P., Van Dyck, M., Verschuuren, M., Kashikar, N. D., Nuydens, R., Timmermans, J.-P., & De Vos, W. H. (2018). Image-Based Profiling of Synaptic Connectivity in Primary Neuronal Cell Culture. Frontiers in Neuroscience, 12. https://doi.org/10.3389/fnins.2018.00389

Vining, K. H., & Mooney, D. J. (2017). Mechanical forces direct stem cell behaviour in development and regeneration. Nature Reviews Molecular Cell Biology, 18(12), 728–742. https://doi.org/10.1038/nrm.2017.108

Vyas, A., Li, Z., Aspalter, M., Feiner, J., Hoke, A., Zhou, C., O’Daly, A., Abdullah, M., Rohde, C., & Brushart, T. M. (2010). An in vitro model of adult mammalian nerve repair. Experimental Neurology. https://doi.org/10.1016/j.expneurol.2009.05.022

Wang, Y., Li, B., Xu, H., Du, S., Liu, T., Ren, J., Zhang, J., Zhang, H., Liu, Y., & Lu, L. (2020). Growth and elongation of axons through mechanical tension mediated by fluorescent-magnetic bifunctional Fe3O4·Rhodamine 6G@PDA superparticles. Journal of Nanobiotechnology. https://doi.org/10.1186/s12951-020-00621-6

Wilson, E. S., & Newell-Litwa, K. (2018). Stem cell models of human synapse development and degeneration. Molecular Biology of the Cell, 29(24), 2913–2921. https://doi.org/10.1091/mbc.E18-04-0222

Yogev, S., Cooper, R., Fetter, R., Horowitz, M., & Shen, K. (2016). Microtubule Organization Determines Axonal Transport Dynamics. Neuron, 92(2), 449–460. https://doi.org/10.1016/j.neuron.2016.09.036

Zhang, S. (2014). Sox2, a key factor in the regulation of pluripotency and neural differentiation. World Journal of Stem Cells, 6(3), 305. https://doi.org/10.4252/wjsc.v6.i3.305

